# Enhancing KCC2 function reduces interictal activity and prevents seizures in temporal lobe epilepsy

**DOI:** 10.1101/2023.09.16.557753

**Authors:** Florian Donneger, Adrien Zanin, Jeremy Besson, Delphine Roussel, Yoness Kadiri, Carla Pagan, Manisha Sinha, Nicolas David, Marion Russeau, Franck Bielle, Bertrand Devaux, Bertrand Mathon, Vincent Navarro, Francine Chassoux, Jean Christophe Poncer

**Affiliations:** Institut du Fer à Moulin, Inserm, Sorbonne Université, UMR-S 1270, 75005, Paris, France; Sorbonne Université, Institut du Cerveau - Paris Brain Institute - ICM, Inserm, CNRS, AP-HP, Hôpital de la Pitié Salpêtrière, 75013, Paris, France; Department of Neuropathology, AP-HP, Hôpitaux Universitaires La Pitié Salpêtrière–Charles Foix, 75013, Paris, France; Department of Neurosurgery, Sainte-Anne Hospital, 75014, Paris, France; AP-HP, Department of Neurosurgery, Pitié-Salpêtrière Hospital, 75013, Paris, France; AP-HP, Epilepsy Unit, Reference Center for Rare Epilepsies, and Department of Clinical Neurophysiology, Pitié-Salpêtrière Hospital, 75013 Paris, France

**Keywords:** chloride transporter, neuropharmacology, hippocampus, drug resistance, resective brain tissue, animal models

## Abstract

The neuronal K/Cl cotransporter KCC2 regulates the transmembrane chloride gradient, which controls the efficacy of GABAergic signaling. In mesial temporal lobe epilepsy (mTLE) and other neurological disorders, reduced KCC2 expression or function can result in depolarizing GABA signaling, which is thought to contribute to pathological activity and seizures. Therefore, restoring chloride homeostasis represents a promising therapeutic strategy. We investigated the mechanisms and antiseizure effects of two small molecules, prochlorperazine (PCPZ) and CLP-257, that have been identified as potential KCC2 enhancers. We found that both compounds enhance KCC2 function and clustering in cortical neurons while reducing its membrane diffusion, without altering canonical regulatory phosphorylation. CLP-257 also selectively increased extrasynaptic, but not synaptic, GABA_A_ receptor-mediated currents. Using in vitro recordings from resected brain tissue of patients with drug-resistant mTLE and in vivo recordings from a mouse model, we show that PCPZ and CLP-257 (or its prodrug CLP-290) effectively suppressed spontaneous epileptiform activity in both models. These findings reveal that PCPZ and CLP-257 act as genuine KCC2 enhancers and provide experimental evidence of the therapeutic potential of such compounds for treating drug-resistant mTLE.

**Significance statement:** A major challenge in treating epilepsy is the high percentage of patients with drug-resistant forms, like mesial temporal lobe epilepsy (mTLE). This study investigates a therapeutic strategy by targeting the neuronal KCC2 transporter, which is often dysfunctional in epilepsy. Our findings identify two compounds, prochlorperazine and CLP-257, that enhance KCC2 function by promoting its clustering on the cell membrane, a previously uncharacterized mechanism. Importantly, these compounds effectively reduce spontaneous epileptiform activity in human brain tissue from mTLE patients and significantly reduce seizure frequency in a mouse model. This work provides a critical proof-of-concept for activating KCC2 as a viable therapeutic approach for drug-resistant epilepsy.

## Introduction

Epilepsy is one of the most common neurological disorders, affecting over 65 million people worldwide (1). Mesial temporal lobe epilepsy (mTLE), the most frequent form of focal epilepsy in adults, is frequently associated with hippocampal sclerosis and often refractory to antiseizure medications (2). Surgical resection of the epileptic focus provides 60 to 75% seizure control at 2 y (3), but not all patients are candidates, and the procedure carries cognitive risks. Moreover, less than 50% of patients remain seizure-free 10 y after surgery (4), highlighting the need for novel therapeutic strategies.

Pharmacoresistant epilepsies, including mTLE, often show altered expression and function of cation-chloride cotransporters (CCCs) that regulate transmembrane chloride gradients (5–10). In neurons, the potassium–chloride cotransporter KCC2 maintains low intracellular chloride, allowing hyperpolarizing inhibition via chloride-permeable GABAA receptors (GABAARs) (11). Reduced KCC2 expression and function, commonly reported in mTLE (5, 9, 12) (but see ref. 13) elevates intraneuronal chloride and can compromise or reverse the driving force for GABAARs (5, 14, 15). Depolarizing and excitatory GABA signaling may then paradoxically enhance neuronal activity and promote the abnormal synchronization underlying epileptiform activities (16–18). Accordingly, pharmacological rescue of chloride homeostasis with bumetanide, an antagonist of the chloride importer NKCC1, suppresses interictal-like activity in human mTLE tissue in vitro (5). Similarly, mutations enhancing KCC2 membrane stability and function (19) or overexpression of recombinant KCC2 (20) reduce acute seizures in animal models. Compensating for CCC dysfunction therefore appears to be a promising therapeutic strategy in pharmacoresistant epilepsy.

However, bumetanide failed to prevent acute neonatal seizures in two clinical trials and induced side effects due to peripheral NKCC1 expression (21, 22). Enhancing the function of the neuron-specific chloride extruder KCC2 is therefore a more selective alternative. Recent high-throughput screens have identified several KCC2 enhancers in heterologous systems (23–25). Among them, the phenothiazine-derived antipsychotic prochlorperazine (PCPZ) and the arylmethylidine family compounds CLP-257 and CLP-290 improved symptoms in animal models of disorders associated with KCC2 downregulation (24, 26–30). Yet, their mechanisms of action remain unknown, and their efficacy and selectivity toward KCC2 is debated (31, 32). In addition, CLP-257 has shown conflicting effects on epileptiform activity in vitro (33, 34), and has mostly been tested in acute seizures such as status epilepticus (35, 36), or phenobarbital-resistant neonatal seizures (29). Only one study tested CLP-290 on epileptogenesis in a chronic epilepsy model, but the reported events lacked clear electrographic features of seizures, raising doubts about efficacy (37). Thus, the impact of KCC2 enhancers on spontaneous chronic seizures in mTLE remains uncertain, and completely unknown in the case of PCPZ.

Here, we first investigated the mode of action and efficacy of PCPZ and CLP-257 on KCC2 in rat hippocampal neurons. Both compounds enhanced KCC2 function, and CLP-257 also acted as a positive allosteric modulator of extrasynaptic but not synaptic GABAARs. Both PCPZ and CLP-257 reduced KCC2 membrane diffusion and increased clustering, without altering total or plasmalemmal KCC2 expression levels, or phosphorylation of canonical regulatory sites. In postoperative human mTLE tissue, both suppressed spontaneous interictal-like discharges. Finally, chronic administration of PCPZ or CLP-290 reduced seizure frequency in a lithium-pilocarpine mouse model of mTLE. Together, our findings reveal the mechanisms and therapeutic potential of two KCC2 enhancers in drug-resistant mTLE.

## Results

### PCPZ and CLP-257 Act as KCC2 Enhancers in Rat Hippocampal Neurons

We first tested whether PCPZ and CLP-257, two candidate KCC2 enhancers (23, 24), potentiate KCC2 in rat hippocampal neurons in vitro. To assess KCC2 function, we measured the somato-dendritic gradient of GABA reversal potential (ΔE_GABA_) using whole cell recordings with a defined intracellular chloride concentration of 29 mM (***Fig. 1***). GABAA receptors were locally activated by Rubi-GABA uncaging at the soma and along the dendrites (50 to 80 µm from the soma) in CLP-257 experiments, or by brief isoguvacine application for PCPZ, as we found that PCPZ was phototoxic to neurons upon exposure to UV light (***Material and Methods*** and ***SI Appendix, Fig. S1***). This assay imposes a controlled chloride load on the neuron, providing a dynamic, quantitative, and specific readout of KCC2-mediated chloride transport. Its specificity is supported by the observation that the gradient is abolished upon KCC2 inhibition but remains unaffected by blockade of the chloride importer NKCC1 (38–40). Compared with gramicidin perforated patch recordings, which only report steady state E_GABA_, the ΔE_GABA_ assay offers greater sensitivity and specificity for detecting changes in KCC2 function.

**Fig. 1.**
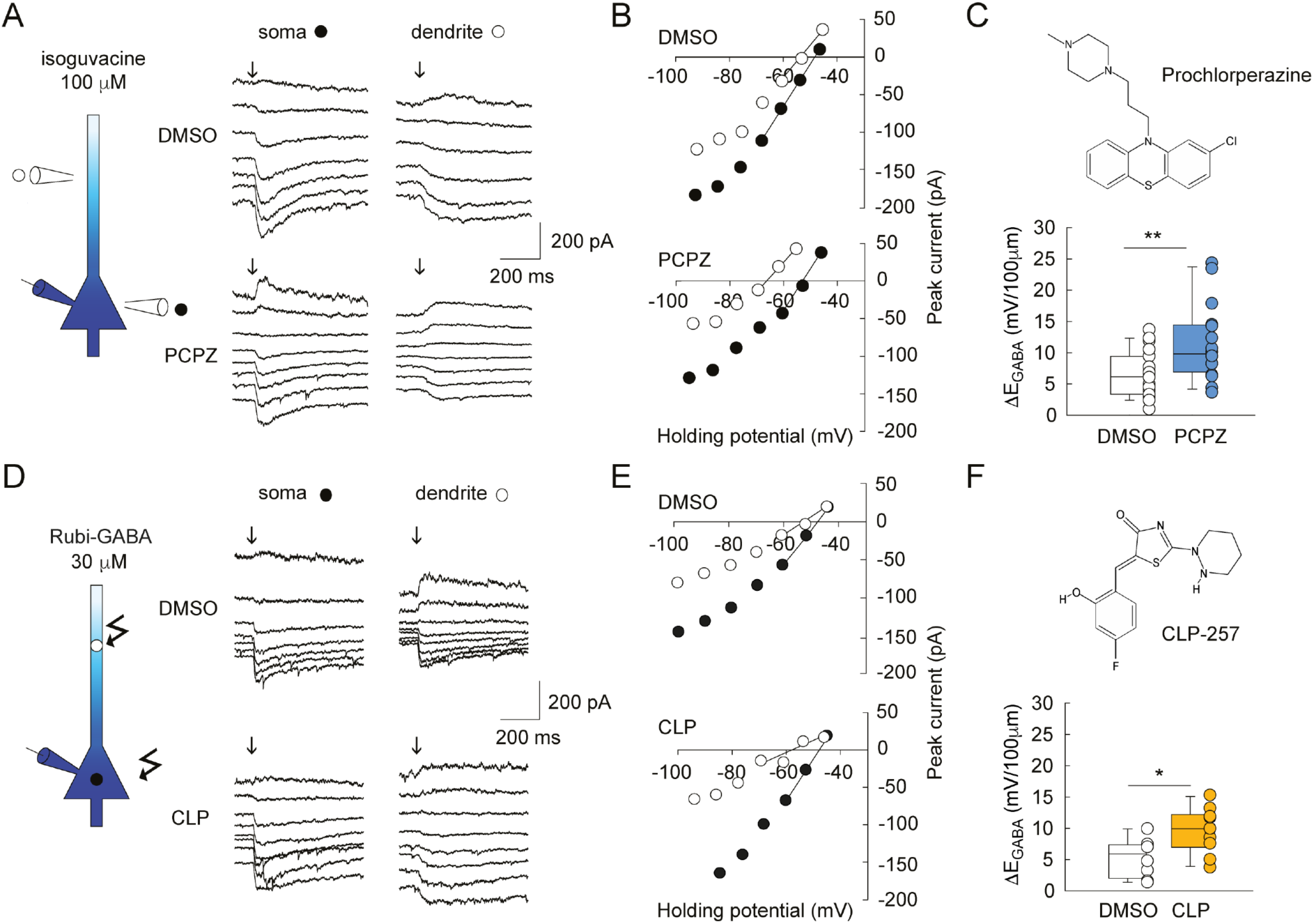
Prochlorperazine (PCPZ) and CLP-257 promote chloride extrusion in hippocampal neurons. (*A*) Representative currents evoked at varying potentials by focal application of isoguvacine (arrow, 100 µM) onto the soma or dendrite of hippocampal neurons (21-24 DIV) previously treated for 2 h with either vehicle only (DMSO, control) or PCPZ (10 µM). (*B*) I-V curves corresponding to the traces shown in *A*. (*C*) 2D chemical structure of PCPZ (*Top*) and summary graph showing somato-dendritic E_GABA_ gradient (*Bottom*, ΔE_GABA_) in control and PCPZ-treated neurons (n = 19 neurons and n = 16 neurons respectively, from five independent cultures). \*\**t* test *P* < 0.01. (*D* and *E*) same as in *A* and *B*, with currents evoked at varying potentials by focal uncaging of Rubi-GABA (arrow, 30 µM) onto the soma or dendrite of hippocampal neurons, pretreated for 2 h with either DMSO alone or 7 µM CLP-257 in DMSO. (*F*) 2D chemical structure of CLP-257 (*Top*) and summary graph showing ΔE_GABA_ (*Bottom*) in control and CLP-treated neurons (n = 8 and 10 neurons, respectively, from four independent cultures) \**t* test *P* < 0.05.

The concentrations of PCPZ (10 µM) and CLP-257 (7 µM) used in these experiments were selected based on effective concentrations reported in previous studies (23, 24). We performed a 2-h treatment with CLP-257 or PCPZ, a paradigm used in previous studies with CLP-257 (23, 41). The mean somato-dendritic ΔEGABA was 6.6 ± 0.8 mV/100 µm in control neurons (DMSO). In neurons preincubated with PCPZ (10 µM), this gradient was significantly increased to 11.5 ± 1.5 mV/100 µm (P = 0.006; Fig. 1 A–C). Similarly, preincubation with CLP-257 (7 µM) increased ΔE_GABA_ from 5.3 ± 1.1 to 9.7 ± 1.1 mV/100 µm (P = 0.014) (***Fig. 1 D–F***). Neither compound had any effect on neuronal membrane resistance or series resistance, or GABAAR conductance (***SI Appendix, Fig. S2***). These data demonstrate that both PCPZ and CLP-257 effectively potentiate KCC2 function in rat hippocampal neurons.

### CLP-257 But Not PCPZ Directly Modulates GABA_A_ Receptor Function

CLP-257 has been suggested to act directly to potentiate GABAARs (31) (but see ref. 32). Because GABAAR-mediated chloride fluxes modulate KCC2 membrane expression and function (42), we asked whether PCPZ and CLP-257 might act primarily to modulate GABAAR function. We tested their effect on GABAAR-mediated currents, using focal application of isoguvacine (100 µM) or uncaging of Rubi-GABA at low concentration (30 µM), respectively (***Fig. 2***). PCPZ had no significant effect on the amplitude (157.5 ± 25.4 vs. 150.7 ± 17.5 pA for control, P = 0.48) or decay time constant (130.8 ± 10.5 vs. 127.7 ± 10.2 ms, P = 0.362) of GABAAR-mediated currents (Fig. 2 A and B). In contrast, CLP-257 rapidly increased their decay time constant (218.2 ± 21.1 vs. 124.4 ± 9.1 ms, P < 0.001) with no detectable effect on their amplitude (162.0 ± 6.6 vs 164.7 ± 4.6 pA for control, P = 0.66; ***Fig. 2 C and D***). This effect of CLP-257 was still observed, albeit weaker, at a 10-fold lower concentration (0.7 µM; 294.1 ± 33.8 ms vs. 239.7 ± 27.2 ms, P < 0.001; ***SI Appendix, Fig. S3***). This concentration is close to the reported EC50 for KCC2 (0.65 µM) (23).

**Fig. 2.**
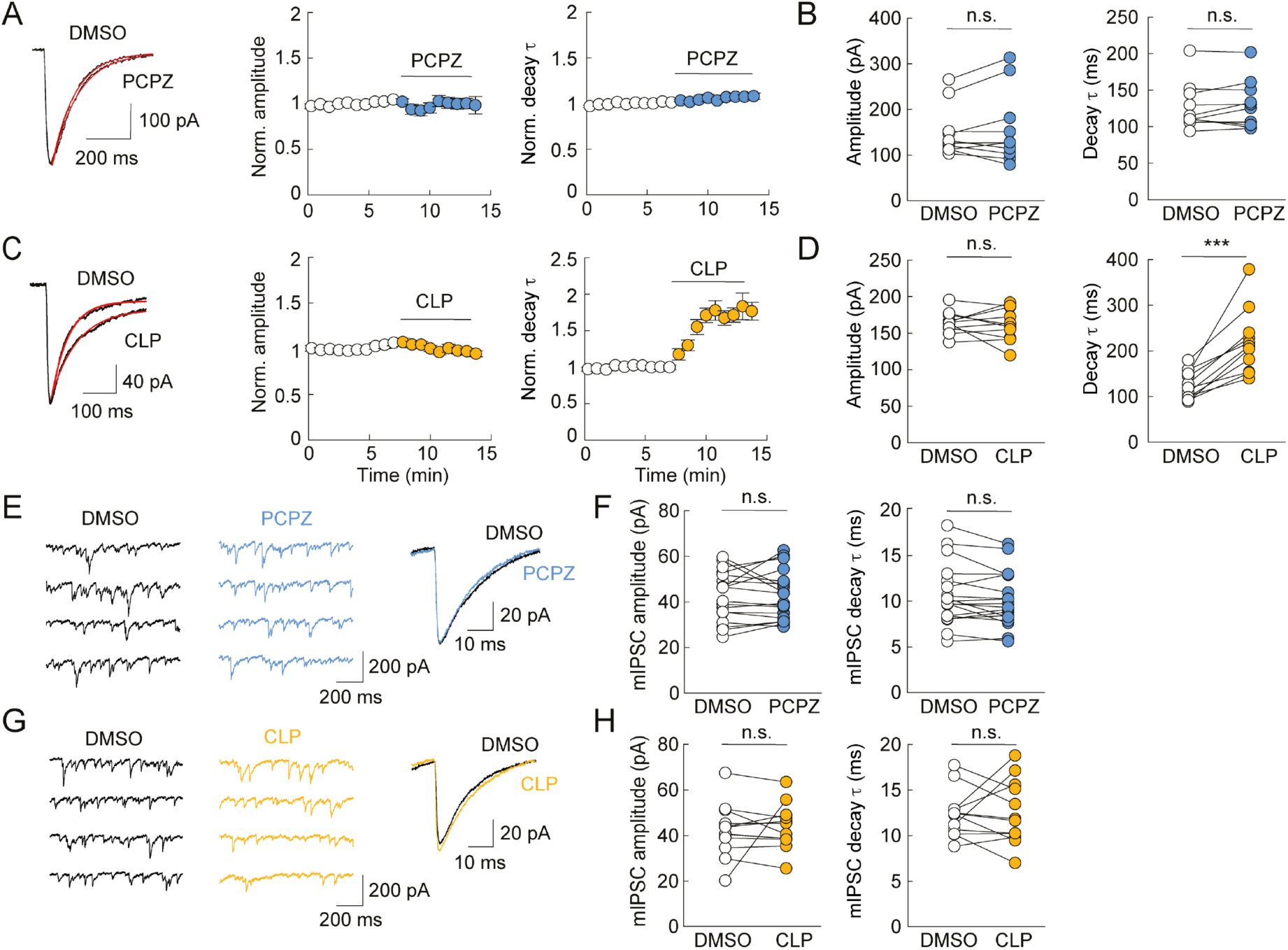
CLP-257 but not PCPZ modulates GABA_A_ receptor function. (*A*, *Left*) Representative recording of currents evoked by focal somatic application of isoguvacine before and during acute application of 10 µM PCPZ. Evoked currents were averaged, and their decay was fit with a simple exponential (red). (*Right*) Normalized amplitude and decay time constant over time of evoked currents. (*B*) Summary graphs showing the effect of PCPZ on the peak amplitude (*Left*) and decay time constant (τ) of isoguvacine-evoked currents (n = 10 neurons each from two independent cultures). (*C* and *D*) Same as in *A* and *B* for currents evoked by Rubi-GABA (30 μM) uncaging before and during application of 7 µM CLP-257 (n = 11 neurons each from two independent cultures. *** Wilcoxon test, *P* < 0.001). (*E*, *Left*) Representative 5-s mIPSC recording before (DMSO) and during application of PCPZ. (*Right*) superimposed averaged mIPSCs detected from this recording. (*F*) Summary graphs showing the effect of PCPZ application on the amplitude (*Left*) and decay time constant (*Right*) of mIPSCs (n = 18 neurons each from four independent cultures. *** Wilcoxon test, *P* < 0.001). (*G* and *H*) Same as *E* and *F* for CLP-257 showing no effect of CLP-257 on mIPSC amplitude and decay time (n = 12 neurons each from four independent cultures. n.s.: not significant).

We next tested the effect of PCPZ and CLP-257 specifically on synaptic GABAARs, by recording miniature inhibitory postsynaptic currents (mIPSCs) from hippocampal neurons. Neither PCPZ nor CLP-257 had detectable effects on mIPSC amplitude (42.9 ± 2.6 vs. 42.5 ± 2.5 pA, P = 0.797 and 43.8 ± 2.9 vs. 42.3 ± 3.4 pA, P = 0.622, respectively) or decay time constant (10.2 ± 0.7 vs. 10.7 ± 0.8 ms, P = 0.086 and 12.5 ± 1.0 vs. 12.4 ± 0.8 ms, P = 0.86, respectively; ***Fig. 2 E–H***). Taken together, these results demonstrate that while both PCPZ and CLP-257 potentiate KCC2 function, CLP-257 also acts to specifically potentiate high-affinity, presumably extrasynaptic but not synaptic GABAAR function in a benzodiazepine-like manner in rat hippocampal neurons.

### PCPZ and CLP-257 Promote KCC2 Clustering and Confinement without Altering KCC2 Expression Levels

We then investigated the mechanisms underlying the potentiation of KCC2 by PCPZ and CLP-257. Enhancement of KCC2 function may result from either increased intrinsic enzymatic activity, increased membrane expression or dimerization of the transporter, or a combination of these factors (43). Therefore, we used surface biotinylation to test whether PCPZ or CLP-257 affects plasmalemmal KCC2 expression. Incubation with PCPZ or CLP-257 for 2 h had no significant effect on total KCC2 expression (***Fig. 3A***, P = 0.137 and P = 0.893, respectively) or on the ratio of surface to total expression (***Fig. 3B***, P = 0.735 and P = 0.8, respectively). In addition, PCPZ or CLP-257 had no significant effect on KCC2 dimerization, as measured by the ratio of KCC2 dimers to monomers (total fraction: P = 0.492 and P = 0.264 respectively, total fraction: P = 0.383 and P = 0.216 respectively; ***SI Appendix, Fig. S4***). Thus, enhanced KCC2 function by PCPZ or CLP-257 does not reflect an increased expression or dimerization of the transporter. PCPZ and CLP-257 also had no effect on total NKCC1 expression (P = 0.102 and P = 0.602, respectively) or on the ratio of surface to total NKCC1 expression (P = 0.266 and P = 0.495, respectively; ***SI Appendix, Fig. S5***).

**Fig. 3.**
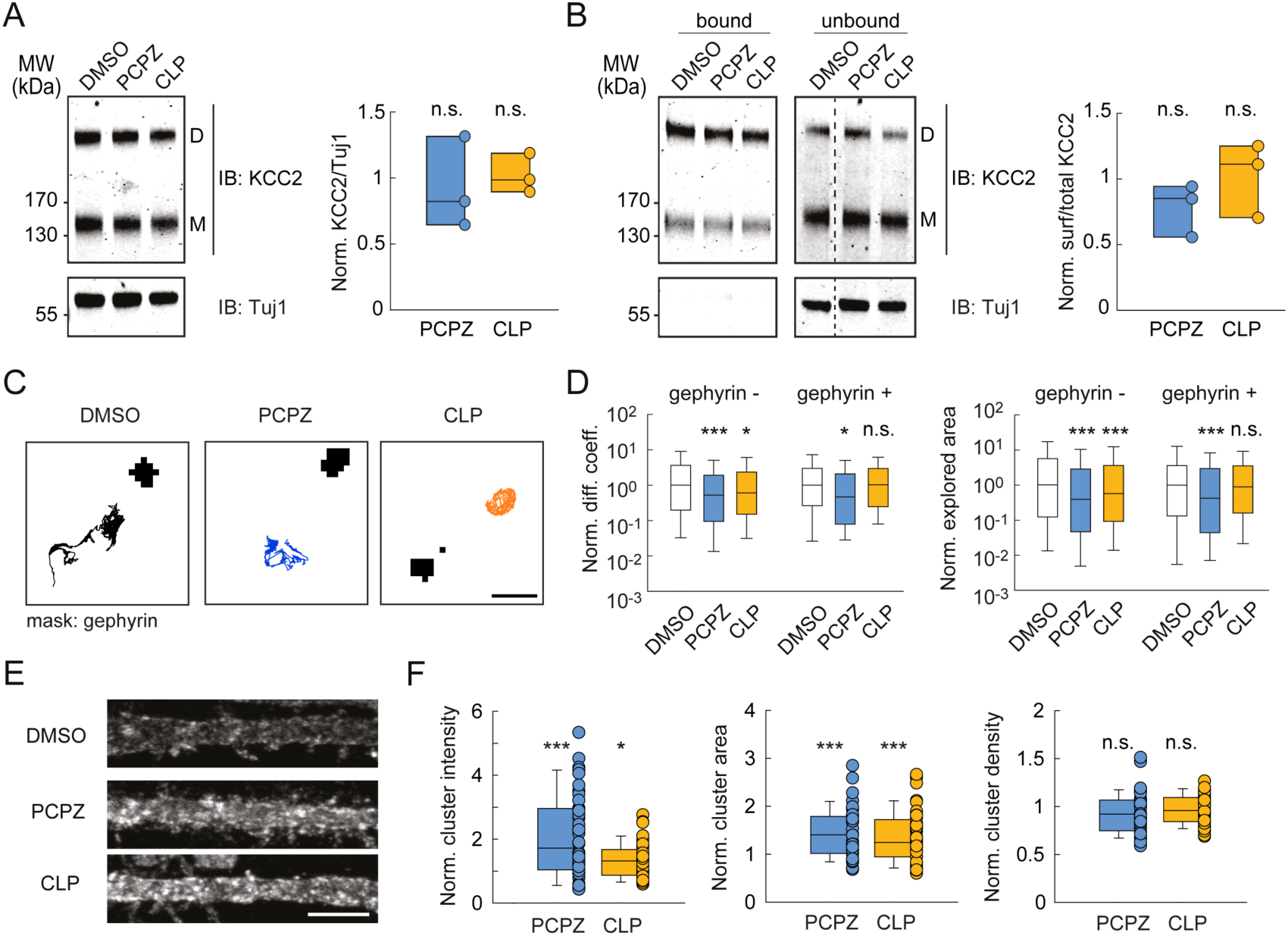
Both PCPZ and CLP-257 reduce KCC2 membrane diffusion and promote its clustering in hippocampal neurons. (*A*, *Left*) representative immunoblots of protein extracts from primary hippocampal cultures treated for 2 h with either PCPZ (10 µM), CLP-257 (7 µM), or DMSO alone. (*Right*) Quantification from three independent experiments. No significant change in total KCC2 expression was detected upon PCPZ or CLP-257 treatment as compared to control (*t* test *P* = 0.14 and *P* = 0.89, respectively). (*B*) Representative immunoblots (*Left*) and quantification (*Right*) showing biotinylated surface KCC2 fraction (bound/total) was also unaffected upon PCPZ or CLP-257 treatment (n = 3 independent cultures, *t* test *P* = 0.74 and 0.80, respectively). (*C*) Representative trajectories of recombinant KCC2 (black) outside inhibitory synapses (gephyrin-FingR mask, gray), as detected using quantum dot-based single particle tracking, showing reduced lateral diffusion upon treatment with PCPZ or CLP-257. Scale, 1 µm. (*D*) Logarithmic distributions of median diffusion coefficients (*D*) and explored areas (EA) of KCC2 trajectories, outside (gephyrin−) or in the vicinity of (gephyrin+) inhibitory synapses, in control neurons (DMSO, D extra n = 614, D syn n = 162 and EA extra n = 1,836, EA syn n = 486 quantum dots) and neurons treated for 2 h with either PCPZ (D extra n = 512, D syn = 131 and EA extra n = 1536, EA syn n = 393 quantum dots) or CLP-257 (D extra n = 697, D syn n = 179 and EA extra n = 2091, EA syn n = 537 quantum dots). Four independent cultures. Mann–Whitney test \**P* < 0.05 \*\*\**P* < 0.001. (*E*) Maximum projections of confocal images of proximal dendrites after KCC2 immunostaining in cultured hippocampal neurons treated for 2 h with PCPZ, CLP-257, or DMSO only. Scale, 5 µm. (*F*) Boxplots showing the distributions of integrated intensity, area and density of KCC2 clusters in control neurons and neurons exposed to either PCPZ or CLP-257. n = 10 neurons per condition per culture and 4 independent cultures per condition. *t* test or Mann–Whitney test \**P* < 0.05 \*\*\**P* < 0.001. n.s.: not significant.

KCC2 function is regulated by fine-tuning of its diffusion and clustering properties at the plasma membrane (44). Thus, the potentiation of KCC2 function is often associated with decreased membrane diffusion and increased clustering (42, 45–47). Membrane lateral diffusion of recombinant Flag-tagged KCC2 was assessed in hippocampal neurons using single particle tracking with quantum dots. The lateral diffusion of KCC2 was reduced by both PCPZ and CLP-257, both in the vicinity of GABAergic synapses and outside. Thus, both compounds reduced the diffusion coefficient of recombinant KCC2 (extrasynaptic −36.5 ± 14.9% P < 0.001; synaptic −26.2 ± 14.0% P = 0.011 for PCPZ and extrasynaptic −20.7 ± 4.9% P = 0.003; synaptic +9.4 ± 11.2% P = 0.97 for CLP-257) and its explored area (extrasynaptic −39.7 ± 17.2% P < 0.001; synaptic −30.6 ± 25.4% P < 0.001 for PCPZ and extrasynaptic −25.1 ± 2.7% P < 0.001; synaptic +4.9 ± 19.4% P = 0.80 for CLP-257; ***Fig. 3 C and D***).

This effect was associated with an increase in KCC2 clustering, as shown by immunocytochemistry assays, which revealed increased cluster integrated intensity (+109.0 ± 61.4% P < 0.001 and +33.9 ± 13.6% P = 0.005, respectively) and area (+45.3 ± 18.1% P < 0.001 and +36.3 ± 20.4% P < 0.001, respectively) with no significant change in cluster density (−6.6 ± 8.4% P = 0.13 and −3.5 ± 4.2% P = 0.34; ***Fig. 3 E and F***). Notably, these effects were not observed after a 30-min application (***SI Appendix, Fig. S6***), suggesting that CLP-257 and PCPZ act slowly to promote KCC2 function. Furthermore, the effect of PCPZ, which is also a D2/D3 receptor antagonist, did not involve dopamine receptor antagonism, since a 2-h treatment with eticlopride produced a small reduction in KCC2 cluster integrated intensity (p = 0.046) and did not significantly affect cluster area or density (p = 0.546 and p = 0.140, ***SI Appendix, Fig. S7***).

Taken together, our results demonstrate that PCPZ and CLP-257 increase net KCC2 function and promote its clustering without affecting total or plasmalemmal KCC2 expression levels. Therefore, PCPZ and CLP-257 likely induce a redistribution of membrane-inserted KCC2, resulting in increased clustering and function.

### PCPZ and CLP-257 Do Not Affect KCC2 Phosphorylation on Canonical Regulatory Residues

The stability, clustering, and membrane turnover of KCC2 are controlled by a variety of posttranslational mechanisms, including phosphorylation of key residues, particularly in its large carboxy-terminal domain, that regulate its net transport function (43, 44, 48). Phosphorylation of serine 940 (S940) by protein kinase C is known to increase the membrane stability and function of KCC2 (45, 49). In contrast, phosphorylation of threonine residues 906 and 1,007 (T906, T1007) by the With-no-lysine (WNK) kinase pathway has been shown to negatively regulate KCC2 function in cortical neurons by affecting its membrane diffusion and clustering (42, 50). We therefore asked whether PCPZ and CLP-257 could act by modulating the phosphorylation of these residues to promote KCC2 function. Immunoprecipitation assays on hippocampo-cortical slices from adult rats (***Fig. 4 A and B*** and ***SI Appendix, Supplementary Methods***) revealed no significant effect of PCPZ or CLP-257 on KCC2 phosphorylation at either S940 (+20.3 ± 18.7%, P = 0.339 and −19.7 ± 34.4%, P = 0.597), T1007 (+26.4 ± 17.0% P = 0.195 and −18.5 ± 17.1%, P = 0.340 respectively), or T906 (−10.2 ± 10.4%, P = 0.384 and −12.7 ± 10.9%, P = 0.308 respectively) residues.

**Fig. 4.**
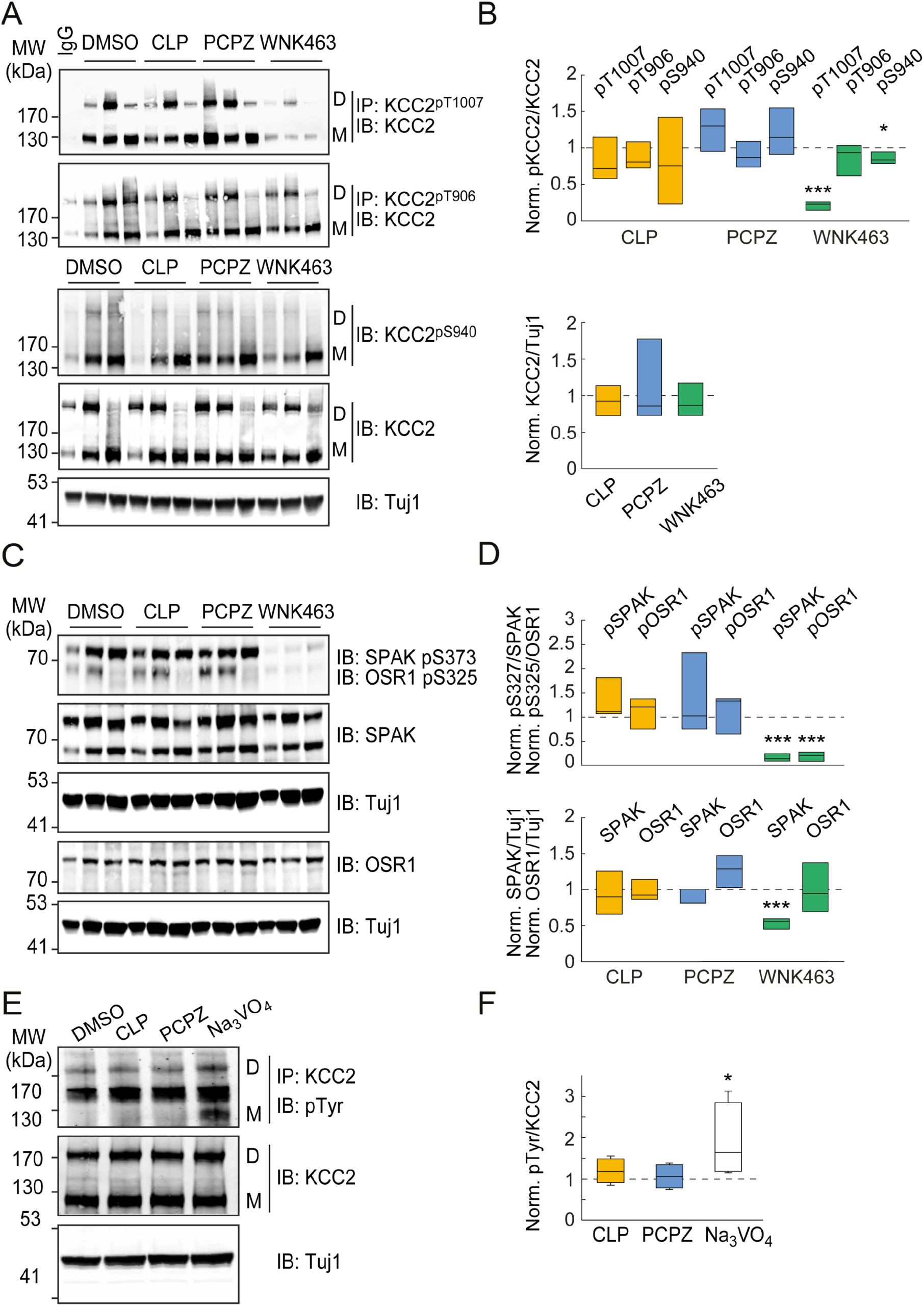
PCPZ and CLP-257 do not affect phosphorylation of canonical KCC2 residues. (*A*) Immunoblots (IB) (*Left*) of protein lysates or proteins immunoprecipitated (IP) with phospho-specific KCC2 antibodies, immunoblotted with antibodies against KCC2 or pS940 KCC2 and beta-tubulin (Tuj1) as control, prepared from hippocampo-cortical slices treated for 2 h with either DMSO alone, CLP-257 (7 µM in DMSO), PCPZ (10 µM in DMSO), or the pan-WNK inhibitor WNK463 (10 µM in DMSO). IgG: control immunoglobulin G. (n = 3 independent replicates 1-3). (*B*) Quantification from the three independent replicates shown in *A*. *t* test \*\*\**P* < 0.001 (*C*) IB (*Left*) of protein lysates, prepared as in *A*, immunoblotted with antibodies against SPAK, OSR1, or their phosphorylated forms and beta-tubulin (Tuj1) as control. (*D*) Quantification from the three independent replicates in *B*. *t* test \*\*\**P* < 0.001. (*E*) Representative IB (*Left*) of proteins immunoprecipitated with an anti-KCC2 antibody and immunoblotted with an anti-phosphotyrosine antibody, prepared from hippocampal-cortices slices treated for 2 h with either DMSO, CLP-257 (7 µM in DMSO), PCPZ (10 µM in DMSO), or sodium orthovanadate (Na_3_VO_4,_ 200 µM). (*F*) Quantification from the four independent replicates shown in *E*. Mann–Whitney test \**P* < 0.05.

Consistent with these results, PCPZ and CLP-257 also had no significant effect on the phosphorylation of the WNK effectors SPAK (S373, +36.9 ± 48.4%, P = 0.488 and +33.3 ± 24.0%, P = 0.238 respectively) and OSR1 (S325, +11.8 ± 23.5%, P = 0.641 and +11.5 ± 18.5%, P = 0.568, respectively) or their total expression levels (SPAK, −12.3 ± 6.4%, P = 0.124 and −6.0 ± 17.4%, P = 0.748, respectively; OSR1, +26.3 ± 12.8%, P = 0.108 and −2.2 ±8.3%, P = 0.800, respectively; ***Fig. 4 C and D***). In contrast, the pan-WNK inhibitor WNK463 (51) reduced the phosphorylation of T1007 (−78.7 ±3.3%, P < 0.001) and to a lesser extent S940 (−14.4 ± 4.7%, P = 0.037) but not T906 (−13.7 ±12.5%, P = 0.333; ***Fig. 4 A and B***). These effects of WNK463 were associated with decreased phosphorylation of SPAK (S373, −84.1 ± 4.6%, P < 0.001) and OSR1 (S325, −81.0 ± 5.6%, P < 0.001; ***Fig. 4 C and D***). Thus, PCPZ and CLP-257 appear to potentiate KCC2 clustering and function independently of detectable changes in phosphorylation of serine 940 and threonine 906 and 1,007 residues. Consistent with this conclusion, these compounds showed no direct inhibitory effect on the activity of WNK1, WNK3, SPAK, or OSR1 in in vitro kinase assays (***SI Appendix, Fig. S8***).

Finally, phosphorylation of residues Y903 and Y1087 has been reported to affect KCC2 clustering and function (47, 52). Therefore, we also tested the effect of PCPZ and CLP-257 on KCC2 tyrosine phosphorylation (***Fig. 4 E and F***). Whereas sodium orthovanadate (Na_3_VO_4_), a tyrosine phosphatase inhibitor, increased KCC2 tyrosine phosphorylation (+89.1 ± 45.4%, P = 0.029), PCPZ, and CLP-257 showed no significant effect (+6.4 ± 14.7% and +19.0 ± 15.0%, P = 1 and P = 0.343, respectively). Taken together, our results demonstrate that PCPZ and CLP-257 potentiate KCC2 function, decrease its diffusion and increase its clustering via a mechanism independent of phosphorylation of canonical regulatory residues in its carboxy-terminal domain.

### PCPZ and CLP-257 Prevent Spontaneous Interictal-Like Discharges in Human mTLE Tissue In Vitro

Human epileptic tissue therapeutically resected from patients diagnosed with mTLE provides an invaluable tool for evaluating antiseizure drug candidates in the pathophysiological context of human mTLE (53, 54). We therefore tested the effect of PCPZ and CLP-257 on 13 tissue samples resected from patients diagnosed with mTLE, 11 of which were associated with hippocampal sclerosis (HS) (***SI Appendix, Table S1***). As previously reported (5, 10), the majority of slices (29/46) prepared from 13 postoperative tissues and recorded with multielectrode arrays (MEA) under control conditions (DMSO, 2 h) showed spontaneous, low-frequency (<40 Hz) events identified as interictal-like discharges (IILDs, ***Fig. 5***). IILDs often originated from only a small subregion of the slice, usually in the subiculum or the dentate gyrus (***Fig. 5 A and B*** and ***SI Appendix, Table S1***). In tissue preincubated with DMSO alone, IILDs had a mean amplitude of 57.3 ± 13.5 µV and occurred at a mean frequency of 0.4 ± 0.1 Hz. As observed previously (5), acute application of the NKCC1 antagonist bumetanide (10 µM) partially reduced the amplitude and frequency of IILDs (***Fig. 5B***). However, these events were almost completely abolished in slices of the same resected tissue incubated for 2 h with either 10 µM PCPZ (***Fig. 5B***) or 7 µM CLP-257 (***Fig. 5C***). Overall, both PCPZ and CLP-257 significantly reduced the proportion of slices with spontaneous IILDs (P < 0.05 for both; ***Fig. 5F***). This effect did not reflect impaired neuronal activity, as IILD and multiunit activity could be restored in slices pretreated with PCPZ or CLP-257, by increasing neuronal excitability with high potassium/low magnesium ACSF (***Fig. 5 D and E***).

**Fig. 5.**
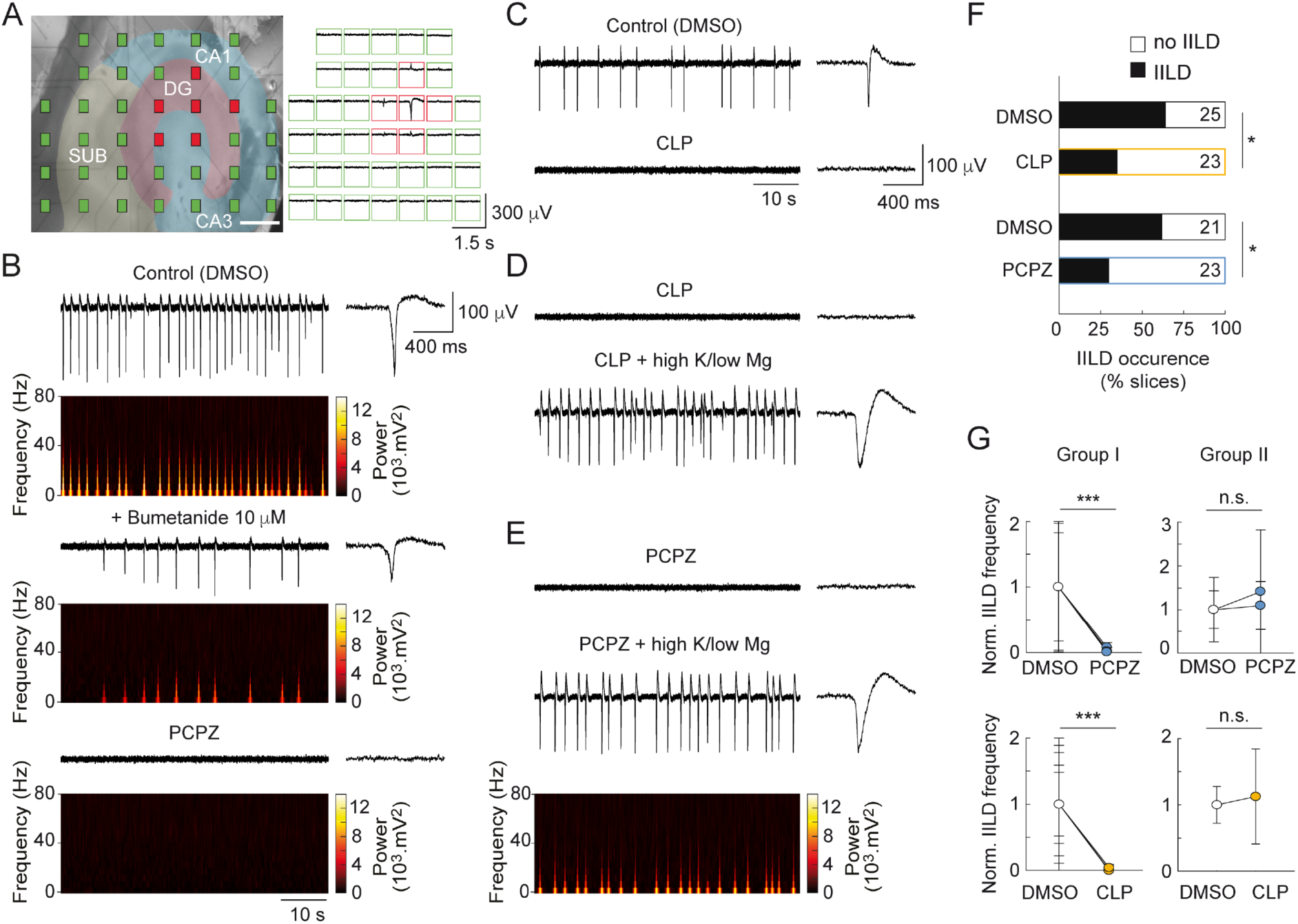
PCPZ and CLP-257 prevent spontaneous interictal-like discharges in temporal lobe slices from mTLE patients. (*A*, *Left*) Photomicrograph of a recorded slice of resected brain tissue from a mTLE patient, showing multielectrode array (MEA) layout. Hippocampal and parahippocampal regions, including the subiculum, were delineated based on slice morphology. Electrodes were color-coded to indicate the type of activity detected (red: interictal-like discharges (IILDs) green: no activity). Scale: 1 mm. (*Right*) Representative recording from this slice, showing synchronous IILDs mostly within the dentate gyrus. (*B*) Representative local field potential recordings and spectrograms showing spontaneous, low-frequency (<40 Hz) IILDs in a slice maintained in control conditions (2 h in DMSO only, *Top*), and their reduction upon >10 min application of 10 µM bumetanide (middle). These events were completely absent from an adjacent slice from the same tissue incubated for 2 h in ACSF supplemented with 10 µM PCPZ (*Bottom*). Insets show individual IILDs. (*C*) Representative recordings showing spontaneous IILDs in a slice from another patient incubated in control conditions (*Top*), and the absence of these events in an adjacent slice kept for 2 h in ACSF supplemented with 7 µM CLP-257 (*Bottom*). (*D* and *E*) Slices from two other patients treated with either CLP or PCPZ also did not display IILDs. However, application of a proconvulsant ACSF (6.5 mM K^+^ and 0.25 mM Mg^2+^) rescued spontaneous IILDs in these slices. (*F*) Summary plot showing the proportion of slices with and without IILDs. *χ* ^2^ test, \**P* < 0.05. (*G*) Graphs showing the mean frequency of spontaneous IILDs (normalized to control) recorded in slices from seven mTLE patients incubated in control ACSF (DMSO) or in the presence of 10 µM PCPZ. Two groups were distinguished: responders (Group I, five patients) and nonresponders (Group II, two patients). Similar data are shown for slices from six patients, incubated in control ACSF or treated with 7 µM CLP-257. Wilcoxon signed rank test, \*\*\**P* < 0.001.

In slices from a group of five patients with mTLE (patients #5, 7, 8, 11, 12; group I; ***SI Appendix, Table S1***), PCPZ virtually abolished IILDs (−97.6 ± 13.8% change in frequency, P < 0.001; ***Fig. 5G***). However, in one slice from patient #3 and in the slices from patient #6 (group II) PCPZ failed to reduce the frequency of IILDs (+25.5 ± 15.9%, P = 0.356; ***Fig. 5G***). Interestingly, neuropathologic examination of the resected tissue from patient #6 revealed an ILAE type 3 HS (***SI Appendix, Table S1***), a rarer pattern of HS often associated with dual pathology (55). Similarly, preincubation with CLP-257 virtually abolished IILDs in slices from 5 out of 6 patients (−96.5 ± 2.3% change in frequency, P < 0.001; ***Fig. 5G***). However, in slices from the remaining patient (#9, ***SI Appendix, Table S1***) CLP-257 failed to reduce the frequency of IILDs (+12.6%, ***Fig. 5G***). Remarkably, neuropathologic examination of this patient’s postoperative tissue did not reveal hippocampal sclerosis, but rather a ganglioglioma associated with a somatic BRAF^V600E^ mutation. Collectively, these data demonstrate that enhancing KCC2 function prevents epileptiform activity recorded in vitro in postoperative tissue from the majority of patients with mTLE and typical ILAE type 1 HS.

### PCPZ and CLP-290 Reduce Interictal Activity and Seizures in a Mouse Model of mTLE

Spontaneous ictal activity is rarely observed in postoperative mTLE tissue slices *in vitro* (53, 54). Therefore, to assess the antiseizure potential of KCC2 enhancers, we tested their effect in a lithium-pilocarpine model of mTLE (56) adapted to mice (***Fig. 6A***). *Status epilepticus* (SE) was induced by intraperitoneal injection of the muscarinic agonist pilocarpine (***SI Appendix, Supplementary Methods***). Mice typically developed SE within 30 to 60 min after pilocarpine injection, and SE was terminated after 1 h by i.p. injection of diazepam and ketamine. ECoG recordings from mice implanted with telemetric probes prior to SE induction revealed intense interictal discharge (IID) activity and occasional seizures for hours to a few days after SE, followed by a seizure-free latent period of 2 to 3 wk before regular spontaneous seizures were detected (***Fig. 6B***). Typical epileptiform activity such as interictal discharges (<20 Hz) or high frequency oscillations [>80 Hz, HFOs, (57–59)] was also observed (***Fig. 6 G and H***). In addition to these electrophysiological biomarkers, typical histological landmarks were detected in the hippocampus of these mice, including hippocampal sclerosis, granule cell dispersion, mossy fiber sprouting, and reduced KCC2 immunoreactivity in the dentate gyrus (***Fig. 6 C and D*** and ***SI Appendix, Fig. S9***). We tested the impact of chronic administration of KCC2 enhancers 5 wk after SE induction (***SI Appendix, Supplementary Methods***). After a 5-d habituation period with twice-daily i.p. vehicle injections (days 35 to 39, control period, 9 AM and 7 PM), we first tested the effect of vehicle vs. PCPZ administration (2 mg/kg, twice daily) on seizure frequency during the next 5 d (days 40 to 44 post-SE, test period) (***Fig. 6 A and E***).

**Fig. 6.**
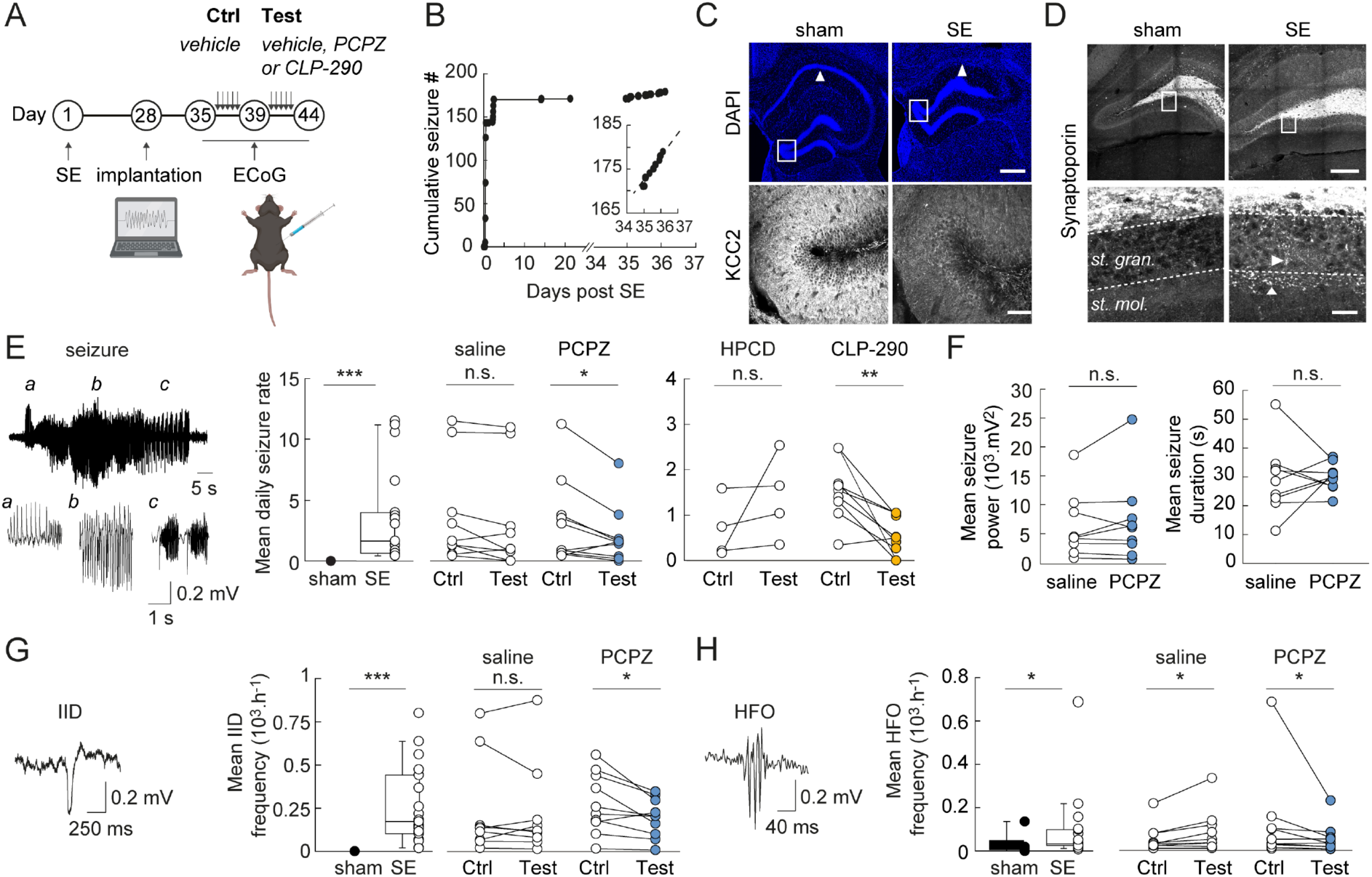
Chronic administration of PCPZ or CLP-290 reduces seizure rate and epileptiform activities in a mTLE mouse model. (*A*) Experimental timeline. (*B*) Graph showing the cumulative number of seizures detected in a mouse immediately after SE and up to 37 d post SE. Recording was interrupted between days 21 and 34 post SE. Note that seizures stop 2 d after SE and chronic seizures occur at a regular rate after day 34 (*Inset*). (*C*, *Top*) Representative confocal fluorescent micrographs of DAPI staining of hippocampal slices from naïve (no SE) and epileptic mice. Arrowheads indicate the CA1 region, which is largely sclerotic in the epileptic mouse. Also note the dispersion of the dentate gyrus granule cells. Scale: 400 µm. (*Bottom*) Confocal fluorescent micrographs of KCC2 immunolabeling in dentate gyrus, from the regions boxed in *Top* panels, showing reduced KCC2 immunofluorescence in the dentate gyrus of the epileptic mouse. Scale: 60 µm (*D*, *Top*) Representative confocal fluorescent micrographs of synaptoporin immunolabeling of the dentate gyrus of control and epileptic mice. Scale: 200 µm. (*Bottom*) Magnification of the regions boxed in *Top* panels, showing ectopic synaptoporin-immunopositive mossy fibers in the stratum granulosum and stratum moleculare only in the epileptic mouse. Scale: 20 µm. (*E*, *Left*) representative ECoG recording of a seizure with *Insets* (a,b,c) showing typical periods of the seizure. Middle, summary plots showing mean seizure frequency per day in sham (no SE, n = 6) and epileptic mice (n = 19). No seizures were detected in sham animals (*** Mann–Whitney test *P* < 0.001). (*Right*) Summary graphs showing mean seizure frequency per hour in epileptic mice receiving i.p. saline injections during the control period (days 35 to 39) and either saline or PCPZ (2 mg/kg in saline) during the test period (days 40 to 44). Mice receiving PCPZ injections during the test period had significantly fewer seizures than during the control period (n = 10 mice, * paired *t* test *P* < 0.05), while no change in seizure frequency was observed in mice receiving saline injections (n = 9, *P* = 0.087). Similarly, mice chronically injected with CLP-290 (100 mg/kg in 20% HPCD) during the test period showed a 55% reduction in mean daily seizure frequency compared to control (n = 9 mice, ** Wilcoxon test *P* < 0.01), while mice receiving vehicle alone (20% HPCD) showed no significant change in seizure frequency (n = 4, *P* = 0.125). (*F*) Summary graphs comparing mean seizure power (*Left*) and duration (*Right*) during the control (saline) and test (PCPZ) periods. While reducing seizure frequency, PCPZ had no significant effect on seizure power (paired *t* test *P* = 0.466) or duration (*P* = 0.657, n = 9 mice). (*G* and *H*, *Left*) Representative ECoG recording of an interictal discharge (IID, in *G*) and a high-frequency oscillation (HFO, in *H*). (*Middle*) Summary plots showing mean IID (in *G*) and HFO (in *H*) frequency in sham (n = 6) and epileptic animals (n = 19). IIDs and HFOs were significantly more frequent in epileptic animals (Mann–Whitney test \**P* < 0.05, \*\*\**P* < 0.001, respectively). Right, summary plots of mean IID (in *G*) and HFO (in *H*) frequency during the control period (days 34 to 39) and the test period (days 40 to 44). Recordings from mice receiving PCPZ injections during the test period displayed significantly reduced IID and HFO frequency as compared to the control period (*paired *t* test *P* < 0.05, n = 10 mice). Recordings from mice receiving saline injections during the test period revealed no change in IID frequency (*P* = 0.846) and a significant increase in HFO frequency (\**P* < 0.05, n = 9 mice).

While seizure frequency was not significantly affected in animals receiving saline during the test period (0.14 ± 0.06 vs. 0.16 ± 0.06 seizures/h, P = 0.087), animals receiving PCPZ showed an approximately 40% reduction in seizure frequency compared to the control period (0.08 ± 0.03 vs. 0.13 ± 0.05 seizures/h, P = 0.032; ***Fig. 6F***). However, PCPZ had no effect on seizure duration (29.9 ± 1.6 vs. 29.1 ± 4.0 s, P = 0.826) or power (7329.9 ± 2414.4 vs. 6451.3 ± 1831.1 mV2, P = 0.329; ***Fig. 6F***), suggesting that it may affect mechanisms involved in seizure onset rather than seizure maintenance. In addition, consistent with our *in vitro* results on postoperative human tissue, PCPZ significantly reduced the frequency of IIDs by approximately 30% compared to control (195.5 ± 35.3 vs. 279.1 ± 55.2 IIDs/h, P = 0.022; ***Fig. 6G***). The effect of PCPZ was even stronger on the frequency of HFOs, with a 55% reduction in PCPZ-treated mice (54.5 ± 21.2 vs. 121.0 ± 64.7 HFO/h, P = 0.014), whereas HFO frequency increased by 55% in the saline-only group (93.3 ± 33.5 vs. 60.3 ± 21.4 HFO/h, P = 0.034; ***Fig. 6H***), showing PCPZ reduces both chronic seizures and interictal activity (including HFOs and IIDs) in an animal model of mTLE. Interestingly, although we observed a circadian rhythmicity in the occurrence of seizures and interictal activity (***SI Appendix, Fig. S10***), as previously reported (60, 61), the effect of PCPZ was similar during light (Zeitgeber time 0 to 12 h) and dark (Zeitgeber time 12 to 24 h) phases (P = 0.551 and P = 0.344 respectively).

Similarly, in another set of experiments, we evaluated the effect of the CLP-257 prodrug CLP-290 (23) on chronic seizures. Mice injected with vehicle alone (20% 2-hydroxypropyl-β-cyclodextrin, HPCD, see ***SI Appendix, Supplementary Methods***) during the control and test periods showed no significant change in seizure frequency (P = 0.13). In contrast, mice treated with CLP-290 (100 mg/kg in 20% HPCD, i.p., twice daily) during the test period showed a 55 % reduction in seizure frequency compared to the control period (P = 0.008; ***Fig. 6E***). Taken together, these results further support that enhancing KCC2 function reduces chronic seizures in an animal model of mTLE.

## Discussion

In this study, we investigated the effect and mode of action of two candidate KCC2 enhancers, PCPZ and CLP-257, in hippocampal neurons. While the concept of restoring chloride gradients for epilepsy is not entirely novel, our study provides critical advancements by revealing a phosphorylation-independent mechanism for KCC2 enhancement by PCPZ and CLP-257. Thus, both compounds reduced KCC2 lateral diffusion and increased its clustering and function, without altering its expression or phosphorylation at canonical regulatory sites. Moreover, we demonstrate the efficacy of both compounds to suppress interictal-like discharges and seizures in human resected brain tissue from drug-resistant mTLE patients as well as in a pertinent animal model of mTLE, thereby emphasizing their direct translational relevance. These findings provide proof-of-concept for the therapeutic potential of KCC2 enhancers in mTLE and potentially other conditions with reduced KCC2 expression.

Several epilepsies are associated with chloride transport dysfunction (62), resulting in paradoxical excitatory signaling through GABAAR (5, 6). Although pro-GABAergic drugs such as benzodiazepines and other GABAAR positive allosteric modulators are often effective in epilepsy (63), their efficacy may be limited when intraneuronal chloride levels are elevated. Restoring chloride gradients therefore appears to be a promising first-line therapeutic strategy in drug-resistant epilepsy.

CL-058 was the first compound identified as a KCC2 enhancer candidate by a library screening in a rodent neuroblastoma-glioma hybrid cell line (23). It was subsequently optimized into CLP-257 and CLP-290, with improved EC_50_ and half-life for in vivo applications. While initially described as specific KCC2 enhancers with little or no activity on other cation-chloride cotransporters, subsequent work found direct binding and potentiating effects on GABAAR (31). Although still controversial (32), these results have called into question CLP drugs as pure KCC2 enhancers. We reconcile these findings by showing that CLP-257 acts both as a bona fide KCC2 enhancer (***Fig. 1***) and as a positive allosteric modulator of extrasynaptic, but not synaptic, GABAAR (***Fig. 2***) in rat hippocampal neurons. These data thus reconcile apparently contradictory results obtained on tonic GABAAR-mediated currents in hippocampal neurons (31) and mIPSCs from spinal cord neurons (32).

PCPZ on the other hand is a first-generation antipsychotic and is generally known as a dopamine D2 antagonist, although it interacts with a lower affinity with serotoninergic, adrenergic, muscarinic, and histaminergic receptors. It was first suggested as a candidate KCC2 enhancer by screening the Prestwick’s library on HEK cells expressing recombinant KCC2 (24) and then shown to reduce the depolarizing driving force of evoked IPSCs in immature rat spinal motoneurons after spinal cord injury. However, its effects on cortical neurons have not been investigated. Our results now show that PCPZ also acts as a potent KCC2 enhancer, with no detectable effect on NKCC1 or GABAAR-mediated currents (***Fig. 2***).

Despite their potential therapeutic interest in restoring chloride transport function in the pathology, the mechanisms of action of CLP-257 and PCPZ on KCC2 have not been fully characterized. Previous studies have suggested that CLP compounds increase total (28, 29) or plasmalemmal KCC2 expression (23, 27, 64) mostly in animal models of chronic pain or spasticity. Similarly, PCPZ has been shown to rescue KCC2 membrane immunoreactivity in lumbar motoneurons after spinal cord injury (24). However, the underlying mechanisms of these effects have not been further investigated. We observed no such changes in either total or plasmalemmal KCC2 expression upon incubation of hippocampal neurons with PCPZ or CLP-257 (***Fig. 3***). Instead, increased KCC2 function was associated with reduced KCC2 membrane diffusion and increased clustering (***Fig. 3***). These observations suggest that under nonpathological conditions, PCPZ and CLP-257 may act by redistributing plasmalemmal KCC2 rather than increasing its abundance.

KCC2 diffusion and clustering in the plasma membrane are tightly correlated with function and are controlled by phosphorylation of key residues in its carboxy-terminal domain (11, 44, 45, 65), making them prime candidate targets for KCC2 modulators. In particular, phosphorylation of S940 and dephosphorylation of T906 and T1007 promote KCC2 clustering and membrane stability (42, 45, 50, 52, 66). This effect usually results in increased membrane expression of the transporter, which was not observed with PCPZ or CLP-257 treatment, making it unlikely that a change in their phosphorylation underlies their effects. Consistent with this conclusion, CLP-257 and PCPZ did not cause a significant change in the phosphorylation of these residues (***Fig. 4***) and did not affect the WNK/SPAK/OSR1 pathway, which is known to phosphorylate T906 and T1007 (42, 66). These observations are consistent with previous observations in a mouse model of ischemia (29). Finally, phosphorylation of Y1087 may be a good candidate target for PCPZ and CLP-257, as it can affect KCC2 clustering and function without affecting its membrane expression (47, 52, 67). However, KCC2 tyrosine phosphorylation was also unaffected by either compound in hippocampo-cortical slices. Further studies will be required to test whether compensatory phosphorylation and dephosphorylation of different tyrosine residues may mask the net effect of KCC2 enhancers on Y1087.

Taken together, our data demonstrate that both PCPZ and CLP-257 enhance KCC2 function and promote its membrane confinement and clustering without altering expression or canonical phosphorylation. Interestingly, other KCC2 enhancers with unknown mechanisms have the same potentiating effect without affecting KCC2 membrane expression or the phosphorylation of S940 or T1007 (36). PCPZ, CLP-257, and probably other KCC2 enhancers may target noncanonical phosphorylation sites, as identified in large-scale proteomics studies (68–70). Alternatively, they may alter KCC2 interaction with various partners known to regulate its function and localization (46, 69, 71), including SNARE proteins (72), CIP1 (73), Neto2 (74), CKB (75), PAM (76), and gephyrin (46). Finally, KCC2 enhancers may also promote KCC2 accumulation in lipid rafts (47), thereby enhancing KCC2 clustering and function.

Although previous studies have reported effects of KCC2 enhancers on epileptiform activity *in vitro* or acute seizures such as *status epilepticus* in animal models, their effects on spontaneous seizures in chronic epilepsy models remain unclear. CLP-257 has been reported to either increase (33) or decrease (34) ictal-like discharges (ILDs) in two *in vitro* models of epileptiform activity. On the other hand, inhibition of the WNK/SPAK/OSR1 pathway with WNK463 prevented KCC2 T1007 phosphorylation and reduced the intensity of kainate-induced status epilepticus in mice (35). Finally, CLP-290 was shown to reduce phenobarbital-resistant neonatal seizures and susceptibility to pentylenetetrazole-induced seizures in a neonatal ischemic mouse model (29), while other KCC2 enhancers prevented the development of kainate-induced *status epilepticus* (36). To date, only one study has investigated the effects of CLP-290 in a chronic epilepsy model (37), focusing solely on early effects on epileptogenesis, and the observed events lacked electrographic features of seizures, raising doubts about their exact nature.

Our findings now show that both PCPZ and CLP-257 not only enhance KCC2 function but also suppress interictal discharges in human hippocampal slices from mTLE patients as well as interictal activity and seizures in a mouse model of mTLE. Notably, out of 13 temporal lobe resections, PCPZ or CLP-257 almost completely suppressed interictal-like activity in 10 samples, whereas it had virtually no effect in three samples (***Fig. 5***). Neuropathologic analysis of these patients revealed that they had rare forms of TLE, one related to a ganglioglioma associated with a somatic mutation of the BRAF proto-oncogene, and the other to ILAE type 3, often associated with a dual pathology. Further investigation of the phenotype of these nonresponding tissues will be necessary to elucidate their pathophysiological characteristics and the origin of their resistance to KCC2 enhancers.

PCPZ is a first-generation FDA-approved antipsychotic with indications in schizophrenia, migraine, and nausea and vomiting (77). Side effects are generally mild in adults but adverse effects such as sedation, extrapyramidal symptoms, and in some rare cases, seizures, have been reported in children (78). Here, we show that PCPZ also has significant antiseizure properties both in postoperative temporal lobe tissue from mTLE patients *in vitro*, and in a mouse model of mTLE. *In vitro*, the effect of PCPZ was comparable to that of bumetanide as described in postoperative epileptic brain tissue from patients with epilepsy of various etiologies (5–7), supporting the idea that restoring chloride homeostasis may be an effective antiseizure strategy. In vivo, PCPZ reduced seizure frequency by approximately 40% (***Fig. 6***) but also greatly reduced interictal discharges and high-frequency oscillations, two hallmarks of epileptic networks (57–59). PCPZ was administered at a concentration of 2 mg/kg by i.p. injection, a dose classically used to test the central effects of PCPZ (79–81). Importantly, the antiseizure effects of PCPZ *in vivo* are unlikely to be due to D2 receptor antagonism, as reduced D2 receptor expression or function is associated with increased, not decreased, seizure threshold and D2 agonists, not antagonists, have been shown to have antiseizure properties (82). This suggests that the KCC2-enhancing property of PCPZ overrides any potential proconvulsant side effects linked to its other pharmacological actions, at least within the experimental conditions tested. In addition, the effects of CLP-290 provide further independent support that promoting KCC2 function reduces seizure frequency in mTLE. Given its dual effect on the function of extrasynaptic GABAAR and KCC2, along with no proconvulsive effect through D2 receptor antagonism, CLP-290 is expected to have a more pronounced antiseizure effect than PCPZ. However, i.p. administration may limit its bioavailability as compared to p.o. administration (23), thereby hindering its antiseizure effect under our experimental conditions.

In conclusion, our results demonstrate that PCPZ and CLP-257 are effective KCC2 enhancers and suggest that KCC2 activation may be an effective strategy in patients with mTLE. Recent progress in elucidating the molecular structure of KCC2 (83–86) provides a unique opportunity to explore the molecular mechanisms underlying the modulation of its expression and function and to design more specific, structure-based KCC2 enhancers with improved pharmacokinetics, efficacy, and specificity.

## Materials and Methods

### Animal Procedures

This study adheres to the ARRIVE guidelines for reporting animal research. All animal procedures were performed in accordance with the European Community Council Directive of November 24, 1986 (86/609/EEC), and the guidelines for the treatment of animals issued by the French Ministry of Agriculture and Forestry (Decree 87-848) and were approved by the “Direction Départementale de la Protection des Populations de Paris” (authorization number D75-05-22). The procedures were evaluated by the local animal welfare structure and the Ethics Committee n°5 (Charles Darwin). Following the positive opinion of the Ethics Committee of our project (APAFIS#24889-202008050923999 v4, APAFIS#5575-2016042916319074 v5, and APAFIS #50258-202407091126447 v3), we obtained the authorization from the Ministry of Research and Higher Education to perform the experiments on animals used for scientific purposes. All procedures were performed in a user facility (either IFM, approved by the Ministry of Agriculture under the reference D-75-05-22, or ICM, reference C-75-13-19).

### Human Tissue and Ethics Approval

Informed consent to participate in the study was obtained from patients or their legal guardians. The protocol was approved by a local research ethics committee (CPP Île-de-France 1 - 14388, January 16, 2017). A small amount of nonessential brain tissue for neuropathologic diagnostic purposes was allocated to research by the neurosurgeon during surgery, in agreement with the neuropathologist and the neurologist. Brain samples were obtained between 2017 and 2021 from 13 patients (four males, nine females, age range: 16 to 63 y, SI Appendix, Table S1) diagnosed with mesial temporal lobe epilepsy that was associated with hippocampal sclerosis (11 patients), glioneuronal tumor (one patient), and hippocampal atrophy with malformation of cortical development (polymicrogyria and subcortical heterotopia, one patient). Patients underwent multimodal preoperative evaluation, including neurological examination, neuropsychological assessment, video-electroencephalographic recording, MRI, and fluorodeoxyglucose positron emission tomography. In addition, stereo-electroencephalography was performed in six patients.

Corticectomies were performed at the Sainte-Anne Hospital (nine patients) and at the Pitié-Salpêtrière Hospital (four patients). Neuropathologic diagnosis was made according to the classification of the Diagnostic Methods Commission of the International League Against Epilepsy (ILAE) (55).

### Statistical Analysis

The number of observations reported corresponds to the number of cells for electrophysiology and immunocytochemistry data, individual quantum dots for single-particle tracking, independent primary hippocampal cultures or animals for biochemistry, animals for in vivo experiments, and slices or patients for human tissue recordings. Sample size selection for experiments was based on published experiments, pilot studies, and in-house expertise. Most experimental analyses were blinded to the treatment conditions. In addition, semiautomated analysis software was used to reduce human bias. Human hippocampal slices were randomly assigned to either the control or treatment paradigm prior to conducting the experiments. Finally, animals were randomly assigned to treatment groups with approximately equal numbers in each condition.

Means are presented ±SEM, medians are presented with their interquartile range (IQR, 25 to 75%). Means were compared using the parametric Student’s t test when statistical conditions (normal distribution and homoscedasticity) were met. Otherwise, the nonparametric Mann–Whitney (MW) test was used. For paired measurements, the Wilcoxon test was used unless statistical conditions for Student’s paired t test were verified. Statistical tests were performed with SigmaPlot 13 (Systat Software) or Prism 10 (GraphPad). Differences were considered significant with p-values less than 5 % (*P ≤ 0.05, **P < 0.01, ***P < 0.001). Error bars correspond to SEM.

A comprehensive description of the experimental and analytical procedures is provided in ***SI Appendix*.**

## Data, Materials, and Software Availability

Software data for MEA and ECoG signal analysis has been deposited on GitHub (https://github.com/jcponcer/PoncerLab/tree/main/Donneger_2026). Some data reported in this paper have been deposited on the public repository Zenodo (https://zenodo.org/records/18668821) within the limits of the data volume accepted by the platform; any additional data exceeding these limits will be shared by the lead contact upon reasonable request.

## Acknowledgments

We wish to thank Liset M de la Prida for helpful discussions and critical reading of the manuscript. We would also like to thank the following people: Jinwei Zhang for his expert guidance with phospho-KCC2 biochemistry; Yves de Koninck and Annie Castonguay for kindly providing CLP-290; Christophe Bernard and Antoine Ghestem for their training on telemetric ECoG probe implantation; Etienne Côme and Marianne Renner for their training and advice on single particle tracking (SPT) recordings and data analysis; Esther Bliard for her assistance with status epilepticus induction; Ludovic Langlois for his assistance during the revision of the manuscript. We are also grateful to the Imaging Facility of the Institut du Fer à Moulin for assistance with confocal imaging, and to the Phenoparc and ePHYS (RRID SCR_026412) facilities at the Paris Brain Institute where the CLP-290 experiments were performed. Some of the data presented in this manuscript was previously described in Florian Donneger’s PhD thesis (https://theses.fr/2022SORUS273). This work was supported by Fondation pour la Recherche Médicale (DEQ20140329539 to J.C.P., FDT202106013286 to F.D.), Agence Nationale de la Recherche ERANET-Neuron (ANR-16-NEU3-0006-02 to J.C.P.), LabEx BioPsy, Agence Nationale de la Recherche (ANR-11-LABX-0035 to J.C.P.), Fondation Française pour la Recherche sur l’Epilepsie (J.C.P.), Federation pour la Recherche sur le Cerveau (J.C.P.), Sorbonne Université (F.D., J.B., and C.P.), and INSERM (J.C.P.). The Poncer lab is affiliated to DIM C-BRAINS, supported by the Conseil Régional d’Ile de France.

## aNotes

Competing interest statement: F.D., A.Z., and J.C.P. are co-inventors on the patent: Methods and Pharmaceutical Compositions for Treating Refractory Epilepsy. WO2022219021, 2022 October.

## References

1 A. K. Ngugi, C. Bottomley, I. Kleinschmidt, J. W. Sander, C. R. Newton, Estimation of the burden of active and life-time epilepsy: A meta-analytic approach. Epilepsia 51, 883–890 (2010).

2 I. Blumcke et al., Histopathological findings in brain tissue obtained during epilepsy surgery. N. Engl. J. Med. 377, 1648–1656 (2017).

3 A. J. Lowe et al., Epilepsy surgery for pathologically proven hippocampal sclerosis provides long-term seizure control and improved quality of life. Epilepsia 45, 237–242 (2004).

4 J. de Tisi et al., The long-term outcome of adult epilepsy surgery, patterns of seizure remission, and relapse: A cohort study. Lancet 378, 1388–1395 (2011).

5 G. Huberfeld et al., Perturbed chloride homeostasis and GABAergic signaling in human temporal lobe epilepsy. J. Neurosci. 27, 9866–9873 (2007).

6 J. Pallud et al., Cortical GABAergic excitation contributes to epileptic activities around human glioma. Sci. Transl. Med. 6, 244ra289 (2014).

7 T. Blauwblomme et al., Gamma-aminobutyric acidergic transmission underlies interictal epileptogenicity in pediatric focal cortical dysplasia. Ann. Neurol. 85, 204–217 (2019).

8 M. Munakata et al., Altered distribution of KCC2 in cortical dysplasia in patients with intractable epilepsy. Epilepsia 48, 837–844 (2007).

9 A. Munoz, P. Mendez, J. DeFelipe, F. J. Alvarez-Leefmans, Cation-chloride cotransporters and GABA-ergic innervation in the human epileptic hippocampus. Epilepsia 48, 663–673 (2007).

10 I. Cohen, V. Navarro, S. Clemenceau, M. Baulac, R. Miles, On the origin of interictal activity in human temporal lobe epilepsy in vitro. Science 298, 1418–1421 (2002).

11 P. Blaesse, M. S. Airaksinen, C. Rivera, K. Kaila, Cation-chloride cotransporters and neuronal function. Neuron 61, 820–838 (2009).

12 H. R. Pathak et al., Disrupted dentate granule cell chloride regulation enhances synaptic excitability during development of temporal lobe epilepsy. J. Neurosci. 27, 14012–14022 (2007).

13 M. R. Karlocai et al., Enhanced expression of potassium-chloride cotransporter KCC2 in human temporal lobe epilepsy. Brain Struct. Funct. 221, 3601–3615 (2016).

14 M. Goutierre et al., KCC2 regulates neuronal excitability and hippocampal activity via interaction with Task-3 channels. Cell Rep. 28, 91–103.e7 (2019).

15 M. R. Kelley et al., Locally reducing KCC2 activity in the hippocampus is sufficient to induce temporal lobe epilepsy. EBioMedicine 32, 62–71 (2018).

16 T. J. Ellender, J. V. Raimondo, A. Irkle, K. P. Lamsa, C. J. Akerman, Excitatory effects of parvalbumin-expressing interneurons maintain hippocampal epileptiform activity via synchronous afterdischarges. J. Neurosci. 34, 15208–15222 (2014).

17 R. Khazipov, G. Valeeva, I. Khalilov, Depolarizing GABA and developmental epilepsies. CNS Neurosci. Ther. 21, 83–91 (2015).

18 G. Di Cristo, P. N. Awad, S. Hamidi, M. Avoli, KCC2, epileptiform synchronization, and epileptic disorders. Prog. Neurobiol. 162, 1–16 (2018).

19 Y. E. Moore, T. Z. Deeb, H. Chadchankar, N. J. Brandon, S. J. Moss, Potentiating KCC2 activity is sufficient to limit the onset and severity of seizures. Proc. Natl. Acad. Sci. U.S.A. 115, 10166–10171 (2018).

20 V. Magloire, J. Cornford, A. Lieb, D. M. Kullmann, I. Pavlov, KCC2 overexpression prevents the paradoxical seizure-promoting action of somatic inhibition. Nat. Commun. 10, 1225 (2019).

21 R. M. Pressler et al., Bumetanide for the treatment of seizures in newborn babies with hypoxic ischaemic encephalopathy (NEMO): An open-label, dose finding, and feasibility phase 1/2 trial. Lancet Neurol. 14, 469–477 (2015).

22 J. S. Soul et al., A pilot randomized, controlled, double-blind trial of bumetanide to treat neonatal seizures. Ann. Neurol. 89, 327–340 (2021).

23 M. Gagnon et al., Chloride extrusion enhancers as novel therapeutics for neurological diseases. Nat. Med. 19, 1524–1528 (2013).

24 S. Liabeuf et al., Prochlorperazine increases KCC2 function and reduces spasticity after spinal cord injury. J. Neurotrauma 34, 3397–3406 (2017).

25 F. J. Prael Iii et al., Discovery of small molecule KCC2 potentiators which attenuate in vitro seizure-like activity in cultured neurons. Front. Cell Dev. Biol. 10, 912812 (2022).

26 A. J. Barbour, K. F. Hauser, A. R. McQuiston, P. E. Knapp, HIV and opiates dysregulate K(+)- Cl(−) cotransporter 2 (KCC2) to cause GABAergic dysfunction in primary human neurons and Tat-transgenic mice. Neurobiol. Dis. 141, 104878 (2020).

27 J. N. Bilchak, K. Yeakle, G. Caron, D. Malloy, M. P. Cote, Enhancing KCC2 activity decreases hyperreflexia and spasticity after chronic spinal cord injury. Exp. Neurol. 338, 113605 (2021).

28 Y. Gao et al., KCC2 receptor upregulation potentiates antinociceptive effect of GABAAR agonist on remifentanil-induced hyperalgesia. Mol. Pain 18, 17448069221082880 (2022).

29 B. J. Sullivan, P. A. Kipnis, B. M. Carter, L. R. Shao, S. D. Kadam, Targeting ischemia-induced KCC2 hypofunction rescues refractory neonatal seizures and mitigates epileptogenesis in a mouse model. Sci. Signal. 14, eabg2648 (2021).

30 I. Keramidis et al., Restoring neuronal chloride extrusion reverses cognitive decline linked to Alzheimer’s disease mutations. Brain 146, 4903–4915 (2023).

31 R. A. Cardarelli et al., The small molecule CLP257 does not modify activity of the K(+)-Cl(−) co-transporter KCC2 but does potentiate GABAA receptor activity. Nat. Med. 23, 1394–1396 (2017).

32 M. Gagnon et al., The small molecule CLP257 does not modify activity of the K(+)-Cl(−) co-transporter KCC2 but does potentiate GABAA receptor activity. Nat. Med. 23, 1396–1398 (2017).

33 S. Hamidi, M. Avoli, KCC2 function modulates in vitro ictogenesis. Neurobiol. Dis. 79, 51–58 (2015).

34 V. I. Dzhala, K. J. Staley, KCC2 chloride transport contributes to the termination of ictal epileptiform activity. eNeuro 8, ENEURO.0208-20.2020 (2020).

35 K. T. Kahle et al., WNK3 modulates transport of Cl- in and out of cells: Implications for control of cell volume and neuronal excitability. Proc. Natl. Acad. Sci. U.S.A. 102, 16783–16788 (2005).

36 R. Jarvis et al., Direct activation of KCC2 arrests benzodiazepine refractory status epilepticus and limits the subsequent neuronal injury in mice. Cell Rep. Med. 4, 100957 (2023).

37 J. Cai et al., The suppressive effect of the specific KCC2 modulator CLP290 on seizure in mice. Epilepsy Res. 203, 107365 (2024).

38 G. Gauvain et al., The neuronal K-Cl cotransporter KCC2 influences postsynaptic AMPA receptor content and lateral diffusion in dendritic spines. Proc. Natl. Acad. Sci. U.S.A. 108, 15474–15479 (2011).

39 Y. Otsu, F. Donneger, E. J. Schwartz, J. C. Poncer, Cation-chloride cotransporters and the polarity of GABA signalling in mouse hippocampal parvalbumin interneurons. J. Physiol. 598, 1865–1880 (2020).

40 S. Khirug et al., GABAergic depolarization of the axon initial segment in cortical principal neurons is caused by the Na-K-2Cl cotransporter NKCC1. J. Neurosci. 28, 4635–4639 (2008).

41 M. Chen et al., APP modulates KCC2 expression and function in hippocampal GABAergic inhibition. Elife 6, e20142 (2017).

42 M. Heubl et al., GABAA receptor dependent synaptic inhibition rapidly tunes KCC2 activity via the Cl−-sensitive WNK1 kinase. Nat. Commun. 8, 1776 (2017).

43 M. A. Virtanen, P. Uvarov, M. Mavrovic, J. C. Poncer, K. Kaila, The multifaceted roles of KCC2 in cortical development. Trends Neurosci. 44, 378–392 (2021).

44 E. Côme, M. Heubl, E. J. Schwartz, J. C. Poncer, S. Lévi, Reciprocal regulation of KCC2 trafficking and synaptic activity. Front. Cell. Neurosci. 13, 48 (2019).

45 I. Chamma et al., Activity-dependent regulation of the K/Cl transporter KCC2 membrane diffusion, clustering, and function in hippocampal neurons. J. Neurosci. 33, 15488–15503 (2013).

46 S. Al Awabdh et al., Gephyrin interacts with the K-Cl cotransporter KCC2 to regulate its surface expression and function in cortical neurons. J. Neurosci. 42, 166–182 (2022).

47 M. Watanabe, H. Wake, A. J. Moorhouse, J. Nabekura, Clustering of neuronal K+-Cl− cotransporters in lipid rafts by tyrosine phosphorylation. J. Biol. Chem. 284, 27980–27988 (2009).

48 K. T. Kahle et al., K-Cl cotransporters, cell volume homeostasis, and neurological disease. Trends Mol. Med. 21, 513–523 (2015).

49 H. H. Lee, T. Z. Deeb, J. A. Walker, P. A. Davies, S. J. Moss, NMDA receptor activity downregulates KCC2 resulting in depolarizing GABAA receptor-mediated currents. Nat. Neurosci. 14, 736–743 (2011).

50 P. de Los Heros et al., The WNK-regulated SPAK/OSR1 kinases directly phosphorylate and inhibit the K+-Cl− co-transporters. Biochem. J. 458, 559–573 (2014).

51 K. Yamada et al., Small-molecule WNK inhibition regulates cardiovascular and renal function. Nat. Chem. Biol. 12, 896–898 (2016).

52 H. H. Lee, R. Jurd, S. J. Moss, Tyrosine phosphorylation regulates the membrane trafficking of the potassium chloride co-transporter KCC2. Mol. Cell. Neurosci. 45, 173–179 (2010).

53 G. Huberfeld et al., Glutamatergic pre-ictal discharges emerge at the transition to seizure in human epilepsy. Nat. Neurosci. 14, 627–634 (2011).

54 C. Le Duigou et al., Imaging pathological activities of human brain tissue in organotypic culture. J. Neurosci. Methods 298, 33–44 (2018).

55 I. Blümcke et al., International consensus classification of hippocampal sclerosis in temporal lobe epilepsy: A Task Force report from the ILAE Commission on Diagnostic Methods. Epilepsia 54, 1315–1329 (2013).

56 G. Foffani, Y. G. Uzcategui, B. Gal, L. Menendez de la Prida, Reduced spike-timing reliability correlates with the emergence of fast ripples in the rat epileptic hippocampus. Neuron 55, 930–941 (2007).

57 L. Menendez de la Prida, A. J. Trevelyan, Cellular mechanisms of high frequency oscillations in epilepsy: On the diverse sources of pathological activities. Epilepsy Res. 97, 308–317 (2011).

58 M. Zijlmans et al., High-frequency oscillations as a new biomarker in epilepsy. Ann. Neurol. 71, 169–178 (2012).

59 M. Levesque, P. Salami, Z. Shiri, M. Avoli, Interictal oscillations and focal epileptic disorders. Eur. J. Neurosci. 48, 2915–2927 (2017), 10.1111/ejn.13628.

60 M. O. Baud, A. Ghestem, J. J. Benoliel, C. Becker, C. Bernard, Endogenous multidien rhythm of epilepsy in rats. Exp. Neurol. 315, 82–87 (2019).

61 P. J. Karoly, et al., Cycles in epilepsy. Nat. Rev. Neurol. 17, 267–284 (2021).

62 R. J. Burman et al., Why won’t it stop? The dynamics of benzodiazepine resistance in status epilepticus. Nat. Rev. Neurol. 18, 428–441 (2022).

63 L. J. Greenfield Jr., Molecular mechanisms of antiseizure drug activity at GABAA receptors. Seizure 22, 589–600 (2013).

64 F. Ferrini, L. E. Lorenzo, A. G. Godin, M. L. Quang, Y. De Koninck, Enhancing KCC2 function counteracts morphine-induced hyperalgesia. Sci. Rep. 7, 3870 (2017).

65 A. M. Hartmann, H. G. Nothwang, NKCC1 and KCC2: Structural insights into phospho-regulation. Front. Mol. Neurosci. 15, 964488 (2022).

66 P. Friedel et al., WNK1-regulated inhibitory phosphorylation of the KCC2 cotransporter maintains the depolarizing action of GABA in immature neurons. Sci. Signal. 8, ra65 (2015).

67 K. Strange, T. D. Singer, R. Morrison, E. Delpire, Dependence of KCC2 K-Cl cotransporter activity on a conserved carboxy terminus tyrosine residue. Am. J. Physiol. Cell Physiol. 279, C860–C867 (2000).

68 A. Cordshagen, W. Busch, M. Winklhofer, H. G. Nothwang, A. M. Hartmann, Phospho-regulation of the intracellular termini of K-Cl cotransporter 2 (KCC2) enables flexible control of its activity. J. Biol. Chem. 293, 16984–16993 (2018).

69 J. L. Smalley et al., Isolation and characterization of multi-protein complexes enriched in the K-Cl co-transporter 2 from brain plasma membranes. Front. Mol. Neurosci. 13, 563091 (2020).

70 M. Weber, A. M. Hartmann, T. Beyer, A. Ripperger, H. G. Nothwang, A novel regulatory locus of phosphorylation in the C terminus of the potassium chloride cotransporter KCC2 that interferes with N-ethylmaleimide or staurosporine-mediated activation. J. Biol. Chem. 289, 18668–18679 (2014).

71 V. Mahadevan et al., Native KCC2 interactome reveals PACSIN1 as a critical regulator of synaptic inhibition. Elife 6, e28270 (2017).

72 H. Asraf et al., SNAP23 regulates KCC2 membrane insertion and activity following mZnR/GPR39 activation in hippocampalneurons. iScience 25, 103751 (2022).

73 M. Wenz, A. M. Hartmann, E. Friauf, H. G. Nothwang, CIP1 is an activator of the K+-Cl− cotransporter KCC2. Biochem. Biophys. Res. Commun. 381, 388–392 (2009).

74 E. A. Ivakine et al., Neto2 is a KCC2 interacting protein required for neuronal Cl− regulation in hippocampal neurons. Proc. Natl. Acad. Sci. U.S.A. 110, 3561–3566 (2013).

75 K. Inoue, J. Yamada, S. Ueno, A. Fukuda, Brain-type creatine kinase activates neuron-specific K+-Cl− co-transporter KCC2. J. Neurochem. 96, 598–608 (2006).

76 N. Garbarini, E. Delpire, The RCC1 domain of protein associated with Myc (PAM) interacts with and regulates KCC2. Cell. Physiol. Biochem. 22, 31–44 (2008).

77 J. S. Furyk, R. A. Meek, D. Egerton-Warburton, Drugs for the treatment of nausea and vomiting in adults in the emergency department setting. Cochrane Database Syst. Rev. 2015, Cd010106 (2015).

78 M. Lau Moon Lin, P. D. Robinson, J. Flank, L. Sung, L. L. Dupuis, The safety of prochlorperazine in children: A systematic review and meta-analysis. Drug Saf. 39, 509–516 (2016).

79 C. Ghelardini, N. Galeotti, C. Uslenghi, I. Grazioli, A. Bartolini, Prochlorperazine induces central antinociception mediated by the muscarinic system. Pharmacol. Res. 50, 351–358 (2004).

80 M. R. Landauer, R. L. Balster, L. S. Harris, Attenuation of cyclophosphamide-induced taste aversions in mice by prochlorperazine, delta 9-tetrahydrocannabinol, nabilone and levonantradol. Pharmacol. Biochem. Behav. 23, 259–266 (1985).

81 C. Ghelardini, N. Galeotti, E. Vivoli, I. Grazioli, C. Uslenghi, The central analgesia induced by antimigraine drugs is independent from Gi proteins: Superiority of a fixed combination of indomethacin, prochlorperazine and caffeine, compared to sumatriptan, in an in vivo model. J. Headache Pain 10, 435–440 (2009).

82 Y. Bozzi, E. Borrelli, The role of dopamine signaling in epileptogenesis. Front. Cell. Neurosci. 7, 157 (2013).

83 X. Chi et al., Cryo-EM structures of the full-length human KCC2 and KCC3 cation-chloride cotransporters. Cell Res. 31, 482–484 (2020).

84 Y. Xie et al., Structures and an activation mechanism of human potassium-chloride cotransporters. Sci. Adv. 6, eabc5883 (2020).

85 T. A. Chew, J. Zhang, L. Feng, High-resolution views and transport mechanisms of the NKCC1 and KCC transporters. J. Mol. Biol. 433, 167056 (2021), 10.1016/j.jmb.2021.167056.

86 S. Zhang et al., The structural basis of function and regulation of neuronal cotransporters NKCC1 and KCC2. Commun. Biol. 4, 226 (2021).

