## Supplementary material for "Enhancing KCC2 function reduces interictal activity and prevents seizures in temporal lobe epilepsy": SI Appendix

**This PDF file includes:**

Supplementary Methods
Supplementary references
Supplementary Table 1
Figures S1 to S10

### Supplementary Methods

#### Primary hippocampal cultures

Primary cultures of hippocampal neurons were prepared as described previously (1-3). Briefly, hippocampi were dissected from embryonic day 18-19 Sprague-Dawley rats of both sexes. Tissue was then trypsinized (0.25% v/v) and mechanically dissociated in 1× HBSS (Invitrogen) containing 10 mM HEPES. Neurons were plated at a density of $120.10^3$ cells/ml on 18-mm diameter glass coverslips precoated with 50 µg/ml poly-D,L-ornithine (Sigma-Aldrich) in plating medium consisting of Minimum Essential Medium (MEM, Sigma-Aldrich) supplemented with horse serum (10% v/v, Invitrogen), L-glutamine (2 mM), and $Na^+$ pyruvate (1 mM) (Invitrogen). After attachment for 3-4 hours, cells were incubated in culture medium consisting of Neurobasal medium supplemented with B27 (1x), L-glutamine (2 mM), and antibiotics (penicillin 200 units/ml, streptomycin, 200 µg/ml; Invitrogen) for up to 4 weeks at 37 °C in a 5% $CO_2$ humidified incubator. One-fifth of the culture medium was replaced each week.

#### Pharmacology

Prochlorperazine dimaleate (Sigma-Aldrich) stock was prepared at 25 mM in dimethyl sulfoxide (DMSO, Sigma-Aldrich) and used at a final concentration of 10 µM. CLP-257 (Tocris) stock was prepared at 7 mM in DMSO and used at a final concentration of 7 or 0.7 µM, as indicated. WNK463 (Tocris) stock was prepared at 50 mM in DMSO and used at a final concentration of 10 µM. Equimolar DMSO concentration was used as a control. Eticlopride hydrochloride stock (Tocris) was prepared at 2 mM in $H_2O$ and used at a final concentration of 2 µM. Sodium orthovanadate (Sigma-Aldrich) stock was prepared at 100 mM in $H_2O$ and used at a final concentration of 100 µM.

For most experiments on hippocampal cultures, neurons were preincubated for 2 hours with either DMSO alone, PCPZ, CLP-257, or eticlopride in a $CO_2$ incubator set at 37°C. Drugs were added directly to the culture medium. For some electrophysiology experiments, neurons were also recorded in the presence of these drugs in the bath. For experiments on rat brain extracts or postoperative human tissue, slices were incubated with either DMSO alone, CLP-257, PCPZ, WNK463 or sodium orthovanadate for 2 hours in interface chambers set at 37°C.

**Cellular electrophysiology**

Neurons were perfused at a rate of 1 ml/min with an extracellular solution containing (in mM): NaCl 120, D-glucose 20, HEPES 10, $MgCl_2$ 3, KCl 2, $CaCl_2$ 2 (pH 7.4), in a recording chamber (Luigs & Neumann) maintained at 33°C and mounted on an upright microscope (BX51WI; Olympus). Neurons were whole-cell patch clamped using borosilicate glass pipettes (Hilgenberg, GmbH) containing (in mM): K-gluconate 104, KCl 25.4, HEPES 10, EGTA 10, MgATP 2, $Na_3$GTP 0.4 and $MgCl_2$ 1.8 (pH 7.4) and held at -65 mV. Recordings were made in the presence of blockers of ionotropic glutamate and $GABA_B$ receptors and voltage-gated sodium channels (in µM): D-APV 50 (HelloBio), NBQX 10 (HelloBio), CGP54626 100 (Tocris), and TTX 1 (Latoxan, France). $GABA_A$R-mediated currents were induced at the somatic or dendritic level (approximately 50-80 µm from the soma) by local uncaging of Rubi-GABA (30 µM, Tocris) using a laser pulse (405 nM, 1-8 ms and 15-80 mW, Omicron Deepstar) delivered through the 40× objective using a photolysis head (Prairie Technologies, Brucker) (4). However, we observed that PCPZ was phototoxic to neurons upon exposure to UV light (**Fig. S1**). Therefore, to test the effect of PCPZ on GABA signaling, currents were therefore evoked by focal application of isoguvacine (100 µM, Sigma-Aldrich) through a second pipette using a PicoSpritzer set at 10 psi for 10 ms (soma) or 100 ms (dendrites). Neurons were voltage clamped from -85 to -5 mV with 3.5

s step increments of 10 mV. The current-voltage relationship of somatic and dendritic currents was then calculated from the peak amplitude of GABAergic currents recorded at each potential, corrected for the liquid junction potential (-15.2 mV in our conditions) and the voltage drop across the series resistance. The reversal potential of $GABA_AR$ currents was estimated for somatic and dendritic activation. The somato-dendritic $E_{GABA}$ gradient ($\Delta E_{GABA}$) was calculated as the difference between somatic and dendritic $E_{GABA}$ normalized to the distance between the two activation sites.

The same protocol was used to test the effect of KCC2 enhancers on extrasynaptic $GABA_A$ receptors, but GABAergic currents were only induced at the somatic level. Neurons were voltage clamped at -65 mV and recorded before (DMSO) and after addition of KCC2 enhancers (PCPZ or CLP-257) in the bath. Series resistance (Rs) and input resistance (Ri) were monitored throughout the recording and cells were discarded if there was a >20% drift in either parameter. $I_{GABA}$ amplitude was calculated as the average peak amplitude of GABAergic currents, recorded during 8 sweeps before and during drug application. To calculate the $GABA_AR$ deactivation time constant, the currents were averaged, and the rate of current deactivation was obtained by fitting their decay to a single exponential.

To test the effect of KCC2 enhancers or eticlopride on synaptic $GABA_A$ receptors, neurons were maintained at 33°C in the same extracellular medium as above and patch-clamped, in whole-cell configuration, with a borosilicate glass pipette containing (in mM): CsCl 135, HEPES 10, EGTA 10, MgATP 4, $Na_3GTP$ 0.4, and $MgCl_2$ 1.8 (pH 7.4) and held at -60mV. Recordings were made in the presence of blockers of ionotropic glutamate receptors and voltage-gated sodium channels (in µM): D-APV 50 (HelloBio), NBQX 10 (HelloBio), and TTX 1 (Latoxan, France). Spontaneous mIPSCs were recorded before (DMSO) and during application of KCC2 enhancers (CLP-257 or PCPZ) or eticlopride in the bath. mIPSC amplitude and frequency were analyzed using Detectivent 4.0. To measure the deactivation time constant of synaptic $GABA_A$Rs, 100 events were averaged, and the rate of current deactivation was obtained by fitting the decay of the averaged currents to a simple exponential.

**Surface biotinylation**

Neurons were washed three times with ice-cold PBS and then incubated with PBS supplemented with 0.5-1mg/ml EZ-Link Sulfo-NHS-SS-Biotin (Pierce, Rockford, IL, USA) at 4°C for 30 minutes with gentle shaking. Biotinylation was stopped by adding Tris-HCl (50 mM; pH 7.4) and cells lysed in a modified RIPA buffer containing (in mM): Tris-HCl 50 (pH 7.4), NaCl 150, 1% Nonidet P-40, 0.5% DOC, 0.1% SDS, NaF 50, $Na_3VO_4$ 1 and protease and phosphatase inhibitors (Roche). Lysates were then collected by gently scraping the bottom of each well. After complete homogenization, the samples were centrifuged, and the supernatant was collected.

A small fraction of the lysates was retained for KCC2 input and total protein quantification using a BCA kit (Thermo Scientific). Lysates were mixed with a 50% slurry of Neutravidin beads (Thermo Scientific) and rotated at 4°C overnight. The beads were then pelleted by centrifugation, and the supernatant (non-biotinylated fraction) was collected. The beads were then washed three times in modified RIPA buffer and once in modified RIPA buffer without detergent. After the last wash, the biotinylated fraction was carefully removed from the beads. Total, biotinylated, and non-biotinylated fractions were then denaturated in 10% β-mercaptoethanol and negatively charged in 6× SDS sample buffer at 37°C for one hour.

Samples were electrophoresed on polyacrylamide gradient gels (4-12%) and transferred to nitrocellulose membranes. The transfer efficiency was checked by the addition of red Ponceau. Membranes were washed with TBS-Tween solution (TBST; Invitrogen) for 30 min and blocked with a solution containing 5% (w/v) skim milk. The membranes were then incubated overnight at 4°C with a rabbit primary antibody against KCC2 (07-432; Millipore; 1:3000) or a goat antibody against NKCC1 (ab99558; abcam; 1:500) and a mouse primary antibody against neuron-specific beta-III tubulin (clone TuJ-1; MAB1195; R&D System; 1:1000). They were then washed three times with a TBST-milk solution and incubated with goat secondary antibodies anti-rabbit DyLight 800 (1:3000; Rockland) or donkey secondary antibodies anti-goat DyLight 800 (1:3000; Rockland), and anti-mouse DyLight 700 (1:1000; Rockland) for one hour at RT. The membranes were then washed again three times with TBST and once with TBS. Fluorescence was detected using an Odyssey infrared imaging system (LI-COR Bioscience). Relative intensities of immunoblot bands were determined by densitometry using ImageJ software. Total KCC2 protein expression was determined as the sum of monomeric and oligomeric bands normalized to beta-III tubulin. Surface expression of KCC2 was determined as the ratio of monomeric + oligomeric biotinylated KCC2 fraction to total KCC2 fraction (biotinylated fraction + non-biotinylated fraction).

**Single particle tracking and analysis**

A previously described KCC2-Flag construct was used for single particle tracking experiments (1). This recombinant Flag-tagged KCC2 transporter was shown to maintain normal trafficking and function in transfected hippocampal neurons.

All constructs were sequenced by Beckman Coulter Genomics (Hope End, Takeley, UK). Neuronal transfections were performed at DIV 13–14 using Transfectin (Bio-Rad), according to the manufacturer's instructions (DNA:transfectin ratio 1 µg:3 µl), with 1-

1.5 μg of plasmid DNA per 20 mm well. Experiments were performed 7–10 days after transfection.

Neurons were stained as described previously (1). Briefly, cells were incubated with a mouse anti-Flag antibody (1:300, Sigma-Aldrich) for 6 min at 37 °C, washed, and incubated with a biotinylated Fab goat anti-mouse antibody (1:300; Jackson Immuno Research) in imaging medium for 6 min at 37 °C. After washing, cells were incubated for 1 min with streptavidin-coated quantum dots (QDs) emitting at 655 nm (1 nM; Invitrogen) supplemented with PBS (1 M; Invitrogen) and 10% casein (v/v) (Sigma). Cells were imaged using an Olympus IX71 inverted microscope equipped with a 60× objective (NA 1.42; Olympus) and a 120 W mercury vapor short arc lamp (X-Cite 120Q, Lumen Dynamics). QD real-time images (30 ms integration time over 1200 consecutive frames) were acquired using an ImagEM EMCCD camera and MetaView software (Meta Imaging 7.7). Cells were imaged within 45 min after pre-incubation with appropriate drugs.

QD tracking and trajectory reconstruction were performed with custom Matlab routines, as described (1, 3). For each imaged neuron, one or two dendritic subregions were quantified. Mean square displacement (MSD) versus time plots were calculated as described and diffusion coefficients (D) were calculated by fitting the first four points of the MSD versus time curves with the equation: $MSD(n\tau) = 4Dn\tau + b$, where b is a constant that reflects the accuracy of spot localization. The explored area of each trajectory was defined as the MSD value of the trajectory at two different time intervals of at 0.42 and 0.45 s.

**Immunocytochemistry**

Neurons were fixed with paraformaldehyde (PFA, 4% w/v in PBS) for 15 min at room temperature (RT). Cells were then washed in PBS, permeabilized with 10% Triton X-100 for 4 min at RT and then exposed to a blocking solution containing goat serum (20% v/v; Invitrogen) and 0.1% Triton X-100 diluted in PBS. Neurons were incubated with rabbit

KCC2 antibody (07-432; Millipore; 1:500) and mouse MAP2 antibody (MAB378; Chemicon; 1:500) for 1 hour at RT. Neurons were then incubated with AlexaFluor 488-conjugated goat anti-mouse antibody (AB_2338840; Jackson Labs; 1:400) and with Cy3-conjugated goat anti-rabbit antibody (111-165-003; Jackson Labs; 1:400) for 45 minutes at RT. Neurons were then labeled with DAPI (1:2000, 5 min), washed in PBS and mounted on slides with Mowiol 844 (48 mg/ml, Sigma).

**Fluorescence image acquisition and analysis**

Images were captured using a 63× objective (NA 1.40) mounted on an upright epifluorescence microscope (DM6000, Leica) equipped with a 12-bit cooled CCD camera (Micromax, Roper Scientific) operated with MetaMorph software (Roper Scientific). For KCC2 cluster detection, the exposure time was adjusted to obtain the best fluorescence-to-noise ratio and to avoid pixel saturation. This exposure time was kept constant across cells and conditions. For experiments on non-transfected cells and for each neuron, dendritic sections were selected using MAP2 labeling and then an image was captured with tetramethylrhodamine (TRITC) filter to visualize KCC2 labeling. KCC2 cluster analysis was performed using MetaMorph software (Roper Scientific), as previously described (5). Images were first flatten-background filtered (kernel size, 3×3×2) to enhance cluster outlines, and a user-defined intensity threshold was applied to select clusters and avoid their coalescence. Clusters were delineated in the area of interest (AOI) and the corresponding regions were superimposed on the raw images to quantify cluster density (mean number of KCC2 clusters per 10 $\mu m^2$), cluster area and integrated cluster intensity (integrated fluorescence pixel intensity within clusters). For each culture, the analysis was performed on 10 cells per condition and approximately 100 clusters per cell.

**Immunoprecipitation and immunoblot**

Male Sprague-Dawley rats, 8-weeks old, were anesthetized by intraperitoneal ketamine/xylazine injection (120/10 mg/kg) and perfused transcardially with ice-cold (0-4°C), oxygenated solution ($O_2$/$CO_2$ 95/5%) containing (in mM): N-methyl-d-glucamine 93, KCl 2.5, $NaH_2PO_4$ 1.2, $NaHCO_3$ 30, HEPES 20, D-glucose 20, ascorbic acid 5, sodium pyruvate 3, $MgSO_4$ 10 and $CaCl_2$ 0.5 (300-310 mOsm, pH7.4). Brains were removed and transverse hippocampal-cortices slices (400 µm thick) were prepared in the same solution using a vibratome (HM650V, Microm). The slices were then incubated for 2 hours with DMSO, KCC2 enhancers (CLP-257, PCPZ), WNK463, or sodium orthovanadate in an interface chamber containing artificial cerebrospinal fluid (ACSF) composed of (in mM): D-glucose 10, KCl 3.5, $NaHCO_3$ 26, $NaH_2PO_4$ 1.25, NaCl 126, $CaCl_2$ 1.6 and $MgCl_2$ 1.2 (290 mOsm), equilibrated with 5% $CO_2$ in 95% $O_2$. After the treatments, the slices were washed with cold PBS, snap frozen with liquid nitrogen, and stored at -80°C. Slices were mechanically lysed on ice, using a 2 ml Dounce, in a buffer containing (in mM): Tris/HCl (pH 7.5) 50, EGTA 1, EDTA 1, sodium orthovanadate 1, sodium-B-glycerophosphate 10, sodium fluoride 50, sodium pyrophosphate 5, sucrose 270, benzamidine 1, phenylmethylsulfonyl fluoride 2, Triton 10%, (v/v) 2-mercaptoethanol 0.1% and protease inhibitors (Roche). After complete homogenization, the samples were centrifuged (16,000 g at 4°C for 10 min) and the supernatant was collected. Protein quantification was performed using the Pierce BCA Protein Assay kit (Thermo Scientific) with BSA as a standard. The following phosphorylation site-specific antibodies: anti-KCC3A phospho Thr1048 (0.35mg/ml, sheep, [S0961C] 1st bleed, Dundee), anti-KCC3A phospho Thr991 (0.26mg/ml, sheep, [S959C] 1st bleed, Dundee) were first incubated with the corresponding non-phosphorylated peptide (10mg/ml, Dundee) at 4°C for 30 min. Phospho-antibodies or anti-KCC2 antibodies (07-432; Millipore) were then coupled with

Protein G–Sepharose (4 Fast Flow; Sigma-Aldrich) at a ratio of 15 µg of antibody per 100 µl beads, for 2 hours at 4°C without agitation. 1.5 mg of clarified cell lysate was incubated with the antibody-coupled bead suspension, overnight at 4 °C with gentle agitation. The supernatant was collected after centrifugation, and the beads were washed twice with NaCl solution (0.25 M NaCl in 1× PBS) and twice with PBS alone. Bound proteins were eluted with 2× LDS sample buffer (Invitrogen) containing 0.5 % (v/v) 2-mercaptoethanol and denatured at 75°C for 20 min (IP with phospho-antibodies) or 37°C for 1 h (IP with anti-KCC2 antibodies) before centrifugation (13,000 g at 4°C for 15-20 min). Samples were subjected to electrophoresis on polyacrylamide gradient gels (4-12% bis-tris gels) and transferred to nitrocellulose membranes. The membranes were rinsed with TBS-Tween solution (TBST; Invitrogen) and blocked with 5% (w/v) skim milk. The membranes were then immunoblotted in 5% (w/v) skim milk in TBST with a primary antibody against KCC2 (rabbit, 07-432; Millipore; 1:1000) or phosphotyrosine (mouse, clone 4G10, Millipore, 1:1000), and a mouse primary antibody against neuron-specific beta-III tubulin (clone TuJ-1; MAB1195; R&D System; 1:1000) overnight at 4 °C. They were then washed three times with distilled water, three times with TBST milk solution, and incubated for 1 hour at RT with goat secondary antibodies anti-rabbit IRDye 800 (1:3000; LI-COR), goat anti-mouse IRDye 700 (1:3000; LI-COR), or donkey anti-sheep DyLight 800 (1:3000; Rockland). The membranes were then washed three times with TBST and once with TBS.

Fluorescence was detected and the relative intensities of immunoblot bands were determined using the Odyssey infrared imaging system (LI-COR Bioscience). For immunoblots without immunoprecipitation, the following antibodies were used for immunodetection of total KCC2, KCC2 phospho S940, total SPAK/OSR1, SPAK phospho Ser373/OSR1 phospho S325: rabbit anti-KCC2 (07-432; Millipore; 1:1000), rabbit anti-KCC2 pS940 (p1551-940, Phosphosolutions), sheep anti-SPAK (S551D, third bleed,

Dundee, 1:1000) and sheep anti-phospho-SPAK (Ser373)/ phospho-OSR1 (Ser325) antibodies (S670B, second bleed, Dundee, 1:1000).

**Kinase profiling assays**

Kinase activity profiling was performed by Eurofins using radiometric assays to measure kinase catalytic activity for PCPZ concentrations ranging from 0.1-500 µM in DMSO or CLP-257 concentrations ranging from 0.1-1000 µM in DMSO, in the presence of 70 µM ATP. Equimolar concentrations of DMSO were used as controls. Additional information on the kinase profiler assay can be obtained from the contractor's website.

**Human tissue and multi-electrode recordings**

Cortical samples obtained in the operating room were immediately transferred to ice-cold (0-4°C), oxygenated solution ($O_2$/$CO_2$ 95/5%) containing (in mM): N-methyl-d-glucamine 93, KCl 2.5, $NaH_2PO_4$ 1.2, $NaHCO_3$ 30, HEPES 20, D-glucose 20, ascorbic acid 5, sodium pyruvate 3, $MgSO_4$ 10 and $CaCl_2$ 0.5 (300-310 mOsm, pH7.4) and transported to the laboratory within 15 to 20 minutes. Transverse hippocampo-subicular slices (400 µm thick) were prepared in the same solution using a vibratome (HM650V, Microm). The slices were maintained at 37°C in an interface chamber containing artificial cerebrospinal fluid (ACSF) consisting of (in mM): D-glucose 10, KCl 3.5, $NaHCO_3$ 26, $NaH_2PO_4$ 1.25, NaCl 126, $CaCl_2$ 1.6 and $MgCl_2$ 1.2 (290 mOsm), equilibrated with 5% $CO_2$ in 95% $O_2$.

For testing the effects of PCPZ and CLP-257, we opted for a two-hour preincubation of the slices to ensure reliable effects on KCC2 clustering and trafficking. This prolonged duration was necessary because our prior observations showed that CLP-257 and PCPZ only induce stable KCC2 changes after prolonged exposure (2 h), not after short applications (30 min)(**Fig. S6**). Testing a range of incubation times was not feasible due to the limited availability of human mTLE tissue.

Multielectrode array recordings were performed using a MEA2100 station (Multi Channel Systems) equipped with a 120-microelectrode array chamber (custom 12x10 layout, 30 µm TiN electrodes, 1 mm vertical and 1.5 mm horizontal spacing). Slices were held in the recording chamber by a homemade platinum-nylon harp and superfused with pre-warmed (37°C) oxygenated ACSF at a rate of 6 ml/min. Slices were imaged using a video microscope stage (MEA-VMT1, MultiChannel Systems) in order to register the position of the electrodes relative to the slice. Extracellular signals were acquired at 10 kHz using the MultiChannel Experimenter (MultiChannel Systems) and analyzed offline using custom software (Matlab, The Mathworks). Semi-automated IILD detection was performed according to a standard procedure (6). Briefly, the signal was denoised, filtered in the 1-40 Hz range, squared and then normalized over the entire recording. IILD detection was then semi-automated using a user-defined threshold.

**Mouse model of temporal lobe epilepsy and ECoG recordings**

8-week-old male C57Bl6/J mice were used in a lithium-pilocarpine based model of temporal lobe epilepsy as described (7). Briefly, animals received an intraperitoneal (i.p.) injection of LiCl (423 mg/kg) and scopolamine (1 mg/kg) 18-24 h and 1 h, respectively, before pilocarpine. Pilocarpine (75 mg/kg) was then injected i.p. and the mice were continuously monitored for the onset of *status epilepticus* (SE). The induction of status epilepticus (SE) was confirmed behaviorally by continuous stage 4-5 seizures according to Racine scale. Mice in which the first injection of pilocarpine did not trigger SE after one hour were given a second injection of 20 mg/kg. One hour after onset, SE was terminated by i.p. injection of diazepam and ketamine (both at 10 mg/kg). All animals received a subcutaneous (s.c.) injection of 500 µl of NaCl 0.9% 1 hour and 4 hours after the termination of SE to minimize dehydration. They were monitored daily for weight and behavior and were provided with enriched liquid chow (Fortimel Energy®) for the first 5-7 days after SE. Mice with >20% weight loss for more than 3 consecutive days were killed and discarded from further analysis.

For chronic PCPZ treatment, control and epileptic mice received two daily i.p. injections of either NaCl 0.9% or PCPZ (2 mg/kg in NaCl 0.9%). For CLP-290 treatment, epileptic mice received two daily i.p. injections of either (2-hydroxypropyl)-β-cyclodextrin (HPCD, 20% in water, pH 4.0) or CLP-290 (100 mg/kg in 20% HPCD). Injections were given 10 hours apart (typically 9 AM and 7 PM).

ECoG probe implantation was performed 28 days after SE in all mice except for 1 mouse that was implanted prior to SE induction in order to monitor chronic seizure onset. Mice were anesthetized with 4.5% isoflurane and maintained on 2-2.5%. For PCPZ experiments, telemetric ECoG probes (model A3028B, Open Source Instruments) were implanted subcutaneously and connected to stainless steel screws (1 mm diameter) stereotaxically inserted in the skull through holes drilled above the right hippocampus (1.94 mm posterior and 1.25 mm lateral with respect to the bregma) and above the cerebellum for reference. Screws were attached to the skull with dental cement. Mice received an s.c. of buprenorphine injection (0.05 mg/kg) 1 hour prior to surgery, an s.c. injection of lidocaine (0.5 mg/kg) immediately prior to the surgery, and two s.c. injections of metacam (5 mg/kg, immediately and 24 hours after the surgery) in order to minimize pain. One week after implantation, the ECoG signal (0.3-160 Hz) was recorded continuously using an Octal telemetric data receiver (Open Source Instruments) and LWDAQ software (Neuroarchiver, Open Source Instruments). The signal was amplified 100-fold and digitized at 512 Hz. Recordings were analyzed offline using custom software (Matlab, The Mathworks). Briefly, the signal was normalized, filtered (bandwidth 5-75 Hz), squared, and an energy threshold-based detection was used to detect seizures (events lasting longer than 5 s). For high frequency oscillations (HFOs) and interictal discharges (IIDs), the ECoG signal was filtered with a minimum order high pass (>80 Hz) and low pass (<20 Hz) filter, respectively, and processed with a Walsh transform. Events were then detected from the transformed signal using a user-defined threshold. Due to the characteristics of our telemetric ECoG recordings, HFO analysis was restricted to frequencies between 80 Hz and 160 Hz, thereby limiting the detection of higher frequency oscillations such as fast ripples (>250 Hz). While HFOs in this range are recognized as biomarkers of epileptiform activity, this frequency band may also encompass physiological ripple activity.

CLP-290 experiments (Fig. 6) were performed on a different recording system. Custom-made enamel-coated stainless-steel ECoG electrodes (#791000; A-M Systems, Sequim, WA) were implanted bilaterally above the hippocampus (1.8 mm posterior and 1.2 mm lateral to the bregma) and the motor cortex (1.8 mm anterior and 1.8 mm lateral to the bregma), and a reference and ground electrode were placed on the cerebellum. Electrode coordinates were derived and adjusted according to the Franklin and Paxinos atlas. After a 1-week recovery period, freely moving implanted mice were connected to an analog-to-digital converter amplifier (Natus Quantum or Brainbox EEG-1166) as part of video ECoG acquisition systems (Neurowork or Deltamed, Natus Medical Incorporated, Pleasanton, CA). EEG signals were recorded at a sampling rate of 2048 Hz and passband filtered between 0.5 and 70 Hz. Simultaneously, video recordings were synchronized with the electrophysiological signals and recorded at 25 frames per second using an Axis night vision camera. Mice were recorded continuously (5 days per week) for up to 3 weeks. One experimenter, blinded to the drug administration protocol, analyzed the ECoG recordings. Seizure detection was performed manually from a sliding window of 20-30 seconds based on specific criteria: abrupt onset and termination of events, amplitude threshold significantly above baseline activity (at least three standard deviations), and/or abnormal changes in background rhythm.

**Immunohistochemistry**

Mice were sacrificed 50 days after SE induction to assess histologic markers of epileptogenesis. Briefly, mice were deeply anesthetized by i.p. injection of Euthasol (180 mg/kg) and transcardially perfused with ice-cold (0-4°C), oxygenated solution ($O_2/CO_2$ 95/5%), containing (in mM): N-methyl-d-glucamine 93, KCl 2.5, $NaH_2PO_4$ 1.2, $NaHCO_3$ 30, HEPES 20, D-glucose 20, ascorbic acid 5, sodium pyruvate 3, $MgSO_4$ 10 and $CaCl_2$ 0.5 (300-310 mOsm, pH 7.4). Brains were removed, fixed in 4% PFA for 14-24 hours and stored in PBS solution containing 30 % sucrose and 0.01 % $NaN_3$. Coronal sections (40 µm thick) were cut with a cryotome (HM450, Microm) at -30°C, collected in PBS and stored at -20°C in Amaral's cryoprotective solution.

Labeling was performed using rabbit antibodies against KCC2 (07-432; Millipore; 1:400) or rabbit anti-synaptoporin (102003; Synaptic System; 1:300) to visualize mossy fibers. After removal from Amaral, the slices were rinsed three times for 1 hour in PBS and then permeabilized for 4 hours in a blocking solution (5 % goat serum and 0.5 % Triton in PBS). Slices were incubated for 48 hours at 4°C with primary antibodies diluted in blocking solution (5 % goat serum and 0.25 % Triton in PBS). Slices were then rinsed three times for one hour in PBS and incubated for 4 hours at RT with donkey anti-rabbit Cy3 secondary antibodies (711-165-152; Jackson Labs; 1:400) diluted in the same blocking solution as above. Slices were rinsed in PB (twice for 15 minutes), labeled with DAPI (1:2000), and incubated with PB overnight. Finally, the slices were mounted with Mowiol. Slices were imaged using a Leica SP5 confocal microscope. The distance between each focal plane was 1 µm and the objective used was 40× (N.A. 1.4).

### Supplementary References

1. I. Chamma *et al.*, Activity-dependent regulation of the K/Cl transporter KCC2 membrane diffusion, clustering, and function in hippocampal neurons. *The Journal of neuroscience : the official journal of the Society for Neuroscience* **33**, 15488-15503 (2013).
2. G. Gauvain *et al.*, The neuronal K-Cl cotransporter KCC2 influences postsynaptic AMPA receptor content and lateral diffusion in dendritic spines. *Proc Natl Acad Sci* **108**, 15474-15479 (2011).
3. M. Heubl *et al.*, GABAA receptor dependent synaptic inhibition rapidly tunes KCC2 activity via the Cl−-sensitive WNK1 kinase. *Nature communications* **8**, 1776 (2017).
4. S. Khirug *et al.*, GABAergic depolarization of the axon initial segment in cortical principal neurons is caused by the Na-K-2Cl cotransporter NKCC1. *The Journal of neuroscience : the official journal of the Society for Neuroscience* **28**, 4635-4639 (2008).
5. S. Al Awabdh *et al.*, Gephyrin Interacts with the K-Cl Cotransporter KCC2 to Regulate Its Surface Expression and Function in Cortical Neurons. *The Journal of neuroscience : the official journal of the Society for Neuroscience* **42**, 166-182 (2022).
6. J. Pallud *et al.*, Cortical GABAergic excitation contributes to epileptic activities around human glioma. *Science translational medicine* **6**, 244ra289 (2014).
7. G. Foffani, Y. G. Uzcategui, B. Gal, L. Menendez de la Prida, Reduced spike-timing reliability correlates with the emergence of fast ripples in the rat epileptic hippocampus. *Neuron* **55**, 930-941 (2007).

| Patient ID | Neuropathology | Sex | Age at surgery | Age at seizure onset (years) | Seizure frequency | SEEG-video | Engel score | Medication | IILD-generating area (*in vitro*) |
|---|---|---|---|---|---|---|---|---|---|
| 1 | Hippocampal sclerosis, ILAE type 1 | F | 43 | childhood (precise date unknown) | monthly | no | IC | LCS, LEV, LTG | undefined |
| 2 | Hippocampal sclerosis, ILAE type 1 | F | 49 | 11 | weekly | no | IA | LCS, LEV | undefined |
| 3 | Hippocampal sclerosis, ILAE type 1 | F | 36 | 23 | monthly | no | IA | LEV, OXC, PRP | DG, Sub. |
| 4 | Unconfirmed diagnosis of hippocampal sclerosis. Atrophy visible on MRI + malformation of cortical development. | F | 37 | 18 | monthly | yes | IV | LTG, ESC, CLB, ESCIT | DG, undefined |
| 5 | Hippocampal sclerosis, ILAE type 1 | F | 63 | 3 | monthly | no | IA | CBZ, GBP, PB | undefined |
| 6 | Hippocampal sclerosis, ILAE type 3 | M | 58 | 40 | monthly | no | ID | CBZ, LCS, CLB, R | DG, Sub. |
| 7 | Hippocampal sclerosis, ILAE type 1 | F | 16 | 12 | monthly | yes | III | LEV, LTG | Sub. only |
| 8 | Hippocampal sclerosis, ILAE type 2 | M | 41 | 8 | monthly | no | IC | LCS, LTG | Sub., undefined |
| 9 | No neuronal loss, ganglioglioma with BRAF mutation | F | 45 | 20 | weekly | yes | IA | TPM, PRP, CLB | DG, Sub. |
| 10 | Hippocampal sclerosis, ILAE type 1 | M | 37 | 29 | weekly | yes | IA | ZNS, CBZ, LTG | undefined |
| 11 | Hippocampal sclerosis, not classifiable | F | 32 | 12 | monthly | yes | IIIA | LTG, PRP, TAC, AZT, EPZ, NVL, PRED, MEL | DG, Sub. |
| 12 | Hippocampal sclerosis, ILAE type 1 | F | 22 | 12 | weekly | no | ID | CBZ, LCS, LTG, PRP | DG only |
| 13 | Hippocampal sclerosis, ILAE type 2 | M | 54 | 12-13 | weekly | yes | IA | CBZ, TPM, ZNS | DG only |

**Supplementary Table 1. Clinical, neuropathological and pharmacological features of the patient cohort.** Abbreviations: AZT: Azathioprine; CBZ: Carbamazepine; CLB: Clobazam; ESCIT: Escitalopram; ESC: Eslicarbazepine; EPZ: Esomeprazole; GBP: Gabapentine; LCS: Lacosamide; LTG: Lamotrigine; LEV: Levetiracetam; MEL: Melatonine; NVL: Nebivolol; OXC: Oxcarbazepine; PRP: Perampanel; PB: Phenobarbital; PRED: Prednisone; R: Ropinirole; TAC: Tacrolimus; TPM: Topimarate; ZNS: Zonesamide. DG: Dentate gyrus. Sub.: subiculum. Grey background indicates patients for which no significant effect of PCPZ or CLP-257 on interictal-like activity was observed *in vitro*.

### Supplementary figures

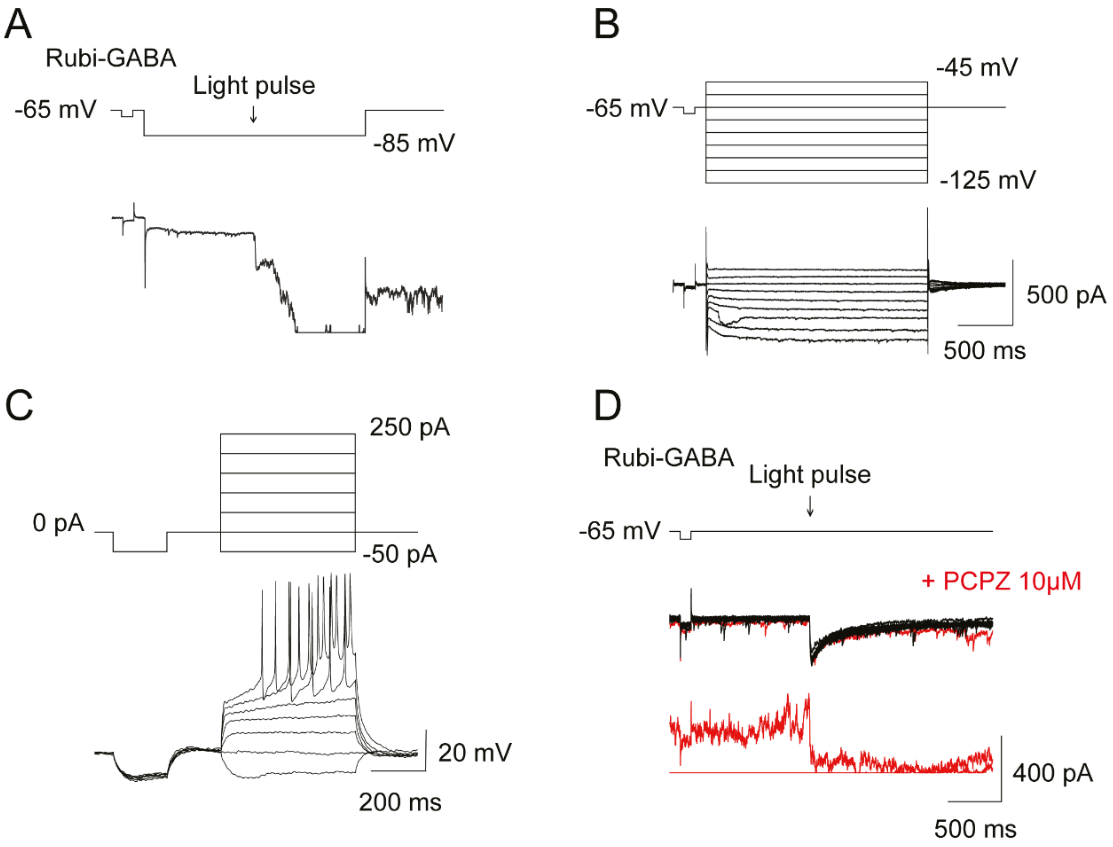


**Figure S1. PCPZ induces UV light phototoxicity in hippocampal neurons *in vitro.* (A)** Representative current evoked at -85mV by focal uncaging of Rubi-GABA (arrow, 30 μM) on the soma of hippocampal neurons, during application of 10 μM PCPZ. The signal is lost after emission of the 405 nm light pulse. (**B)** Representative voltage-clamp recordings of a neuron at different potentials (from -125 to -45 mV), in the presence of PCPZ (10 μM). The recording is stable over time in the absence of the light pulse. (**C)** Representative current-clamp recordings from the same neuron as in B. A normal firing pattern is observed after the application of PCPZ. (**D)** Superimposition of 34 representative traces of GABAergic currents induced by photolysis of Rubi-GABA (arrow) before (black) and after (red) PCPZ application. The addition of PCPZ is associated with signal loss, suggesting a phototoxicity of this compound in blue light (405nm).

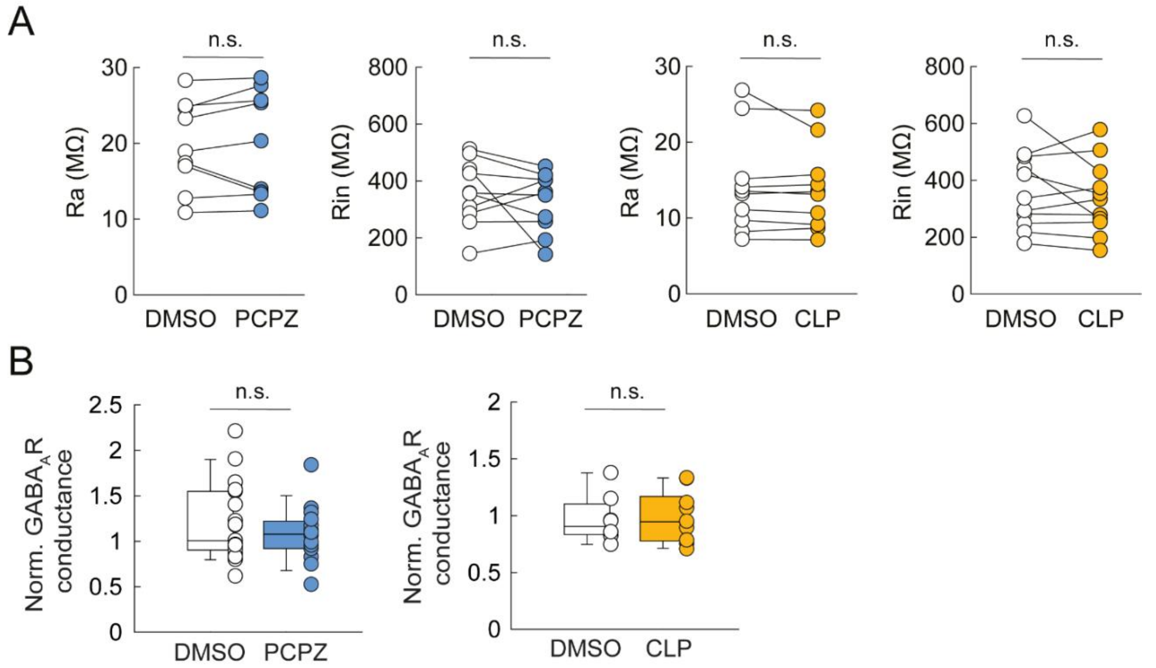


**Figure S2. PCPZ and CLP-257 do not affect membrane properties or $GABA_A$ receptor conductance. (A)** Summary plots showing the lack of significant effect of PCPZ or CLP-257 application on series (Ra) or membrane (Rin) resistance (n=10 and n=11 neurons, respectively, from 2 independent cultures. Wilcoxon test, n.s.: not significant). **(B)** Summary graph showing normalized $GABA_A$R conductance in control vs PCPZ (n=19 and n=16 neurons, respectively, from 5 independent cultures) or control vs. CLP-257-treated neurons (n=8 and n=10 neurons, respectively, from 4 independent cultures. Wilcoxon test, n.s.: not significant).

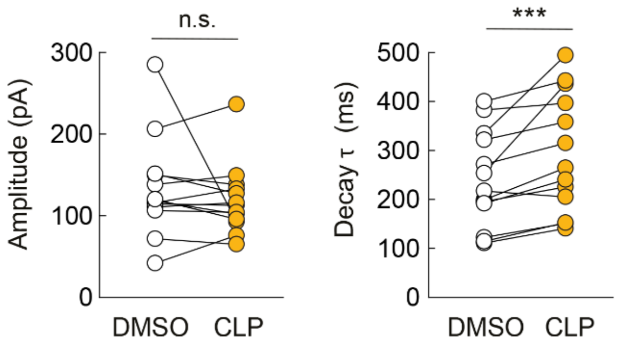


**Figure S3. CLP-257 modulates $GABA_A$ receptor function at submicromolar concentrations.** Summary plots showing the effect of 0.7 μM CLP-257 on the peak amplitude (left) and decay time constant (right) of currents evoked by focal uncaging of Rubi-GABA (30 μM) on hippocampal neurons in vitro (n=13 neurons from 3 independent cultures each. *** Wilcoxon test, $p<0.001$, n.s.: not significant).

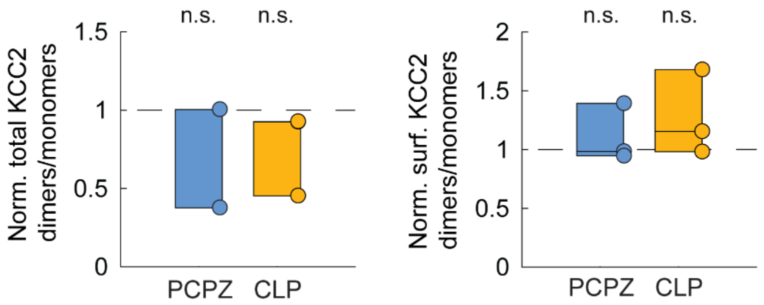


**Figure S4. PCPZ and CLP-257 do not affect KCC2 dimerization.** Summary graphs showing the fractions (normalized to control) of total (left) and surface (right) KCC2 dimers to monomers. No significant effect was observed after treatment with PCPZ or CLP-257 (n=3 independent cultures. t-test, n.s.: not significant).

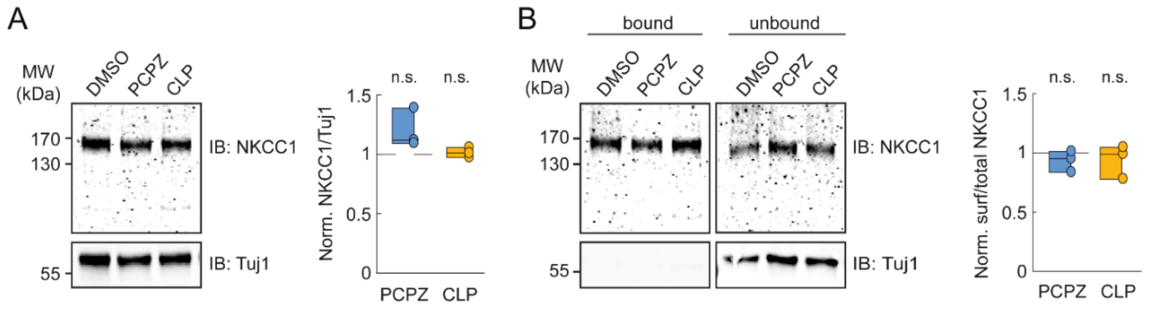


**Figure S5. PCPZ and CLP-257 do not affect total or plasmalemmal expression levels of NKCC1. (A)** Representative immunoblots (left) of protein extracts from hippocampal cultures incubated for 2 hours with either PCPZ (10 μM in DMSO), CLP-257 (7 μM in DMSO), or DMSO alone. Right, quantification of 3 independent experiments. No significant change in total NKCC1 expression was detected upon PCPZ or CLP-257 treatment as compared to control (t-test, n.s.: not significant). (**B)** Immunoblots (left) and quantification (right) showing that the biotinylated surface NKCC1 fraction (bound/total) was also unaffected by incubation with PCPZ or CLP-257 (n=3 independent cultures. t-test, n.s.: not significant).

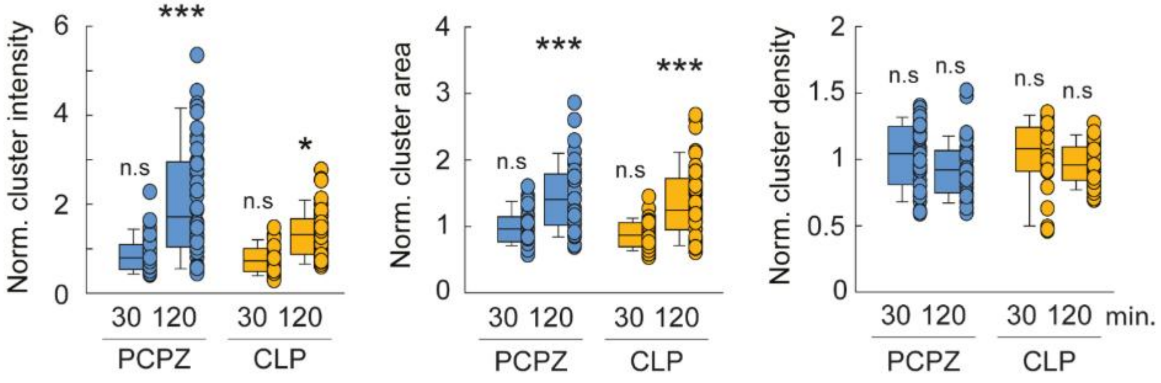


**Figure S6. Time dependence of PCPZ- and CLP-257- induced changes in KCC2 clustering.** Boxplots showing the distributions of the integrated intensity, area and density of KCC2 clusters in control neurons and in neurons exposed to either PCPZ or CLP-257, for 30 minutes or 2 hours. Note that significant effects of both drugs are observed only after 2 hours of treatment, while no significant changes are detected after 30 minutes. n = 10 neurons per condition per culture, 4 independent cultures per condition, except for CLP-257 at 2 hours (3 independent cultures). Statistical significance was assessed using t-test or Mann–Whitney test: $*p < 0.05$, $**p < 0.01$, $***p < 0.001$.

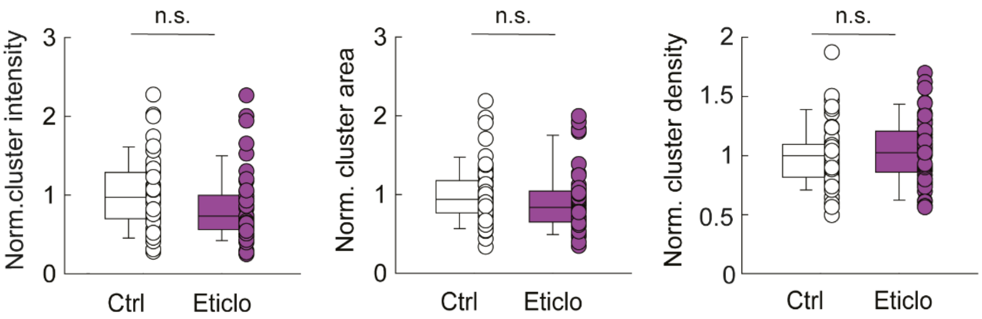


**Figure S7. The D2R antagonist eticlopride does not affect KCC2 clustering in hippocampal neurons.** Box plots showing the distribution of normalized integrated intensity (left), area (middle), and density (right) of KCC2 clusters in control neurons and neurons incubated with 2 µM eticlopride for 2 hours. No significant difference was found between control and eticlopride-treated neurons (n=10 neurons per condition per culture, and 5 independent cultures per condition, Mann-Whitney test, n.s.: not significant).

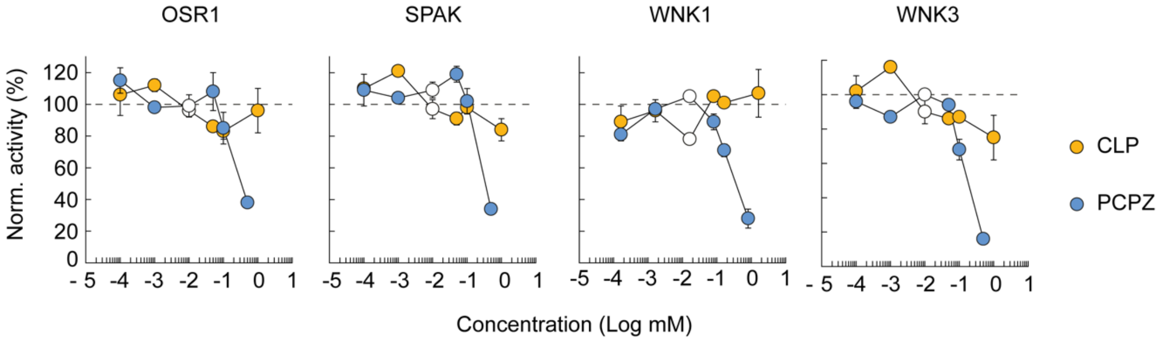


**Figure S8. PCPZ and CLP-257 do not inhibit WNK/SPAK/OSR1 kinase activity.** Kinase activity profile of recombinant human OSR1, STK39 (SPAK), WNK1 and WNK3 in the presence of PCPZ (blue symbols) or CLP-257 (orange symbols) concentrations ranging from 0.1-500 and 1-1000 µM, respectively. Means±SD of 2 replicates were normalized to control (DMSO). Open symbols represent the closest concentration of PCPZ and CLP-257 used in all other in vitro assays in our study (10 and 7 µM, respectively). No detectable inhibition of any kinase was observed at these concentrations.

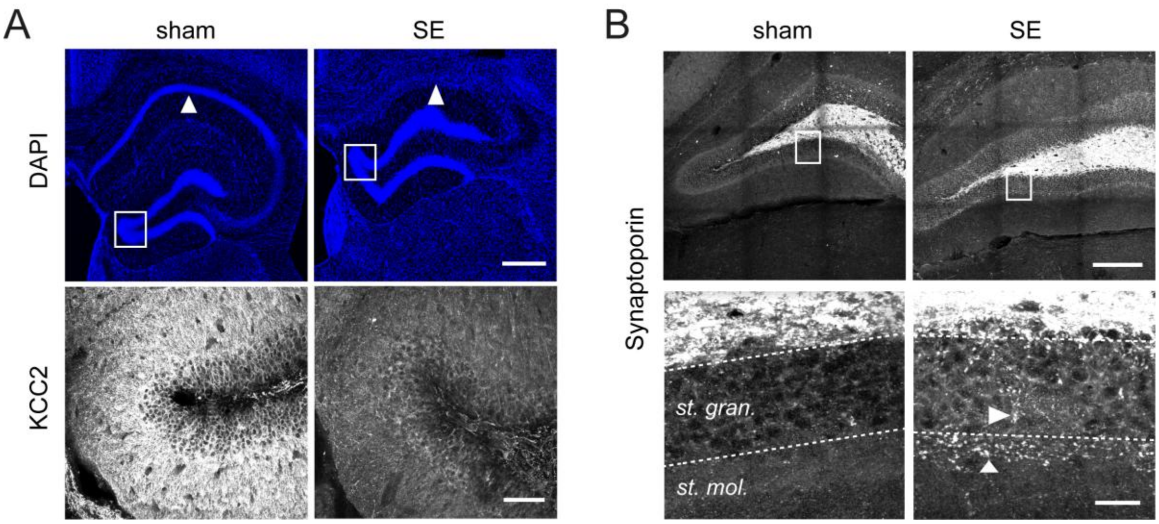


**Figure S9. Lithium-pilocarpine epileptic mice display characteristic hippocampal histological features of mesial temporal epilepsy with hippocampal sclerosis** (enlarged illustrations from Figure 6). **(A)** Top, representative confocal fluorescent micrographs of DAPI staining of hippocampal slices from naïve (no SE) and epileptic mice. Arrowheads indicate the CA1 region, which is largely sclerotic in the epileptic mouse. Also note the dispersion of the dentate gyrus granule cells. Scale: 400 µm. Bottom, confocal fluorescent micrographs of KCC2 immunolabelling in dentate gyrus, from the regions boxed in top panels, showing reduced KCC2 immunofluorescence in the dentate gyrus of the epileptic mouse. Scale: 60 µm **(B)** Top, representative confocal fluorescent micrographs of synaptoporin immunolabelling of the dentate gyrus of control and epileptic mice. Scale: 200 µm. Bottom, magnification of the regions boxed in top panels, showing ectopic synaptoporin-immunopositive mossy fibers in the stratum granulosum and stratum moleculare only in the epileptic mouse. Scale: 20 µm.

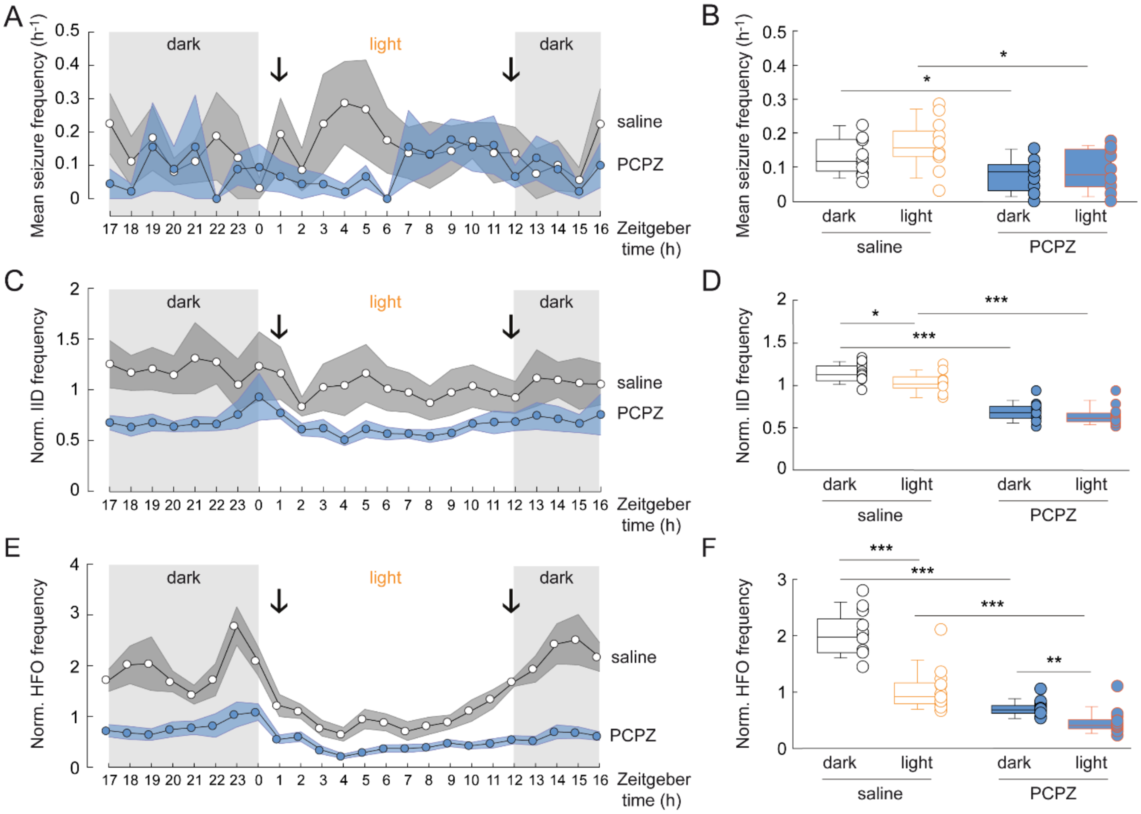


**Figure S10. PCPZ reduces seizure frequency and interictal activity independent of their circadian modulation. (A)** Summary plot showing averaged seizure frequency during the day (number of seizures per hour/number of recording days, mean±SEM) during the test period (days 40 to 44 post-SE), for mice injected twice daily (black arrows) with saline (gray, n=8) or PCPZ (2 mg/kg, blue, n=9). Light and dark periods are indicated. The time scale is the Zeitgeber time. ZT0=time when light is turned on (8 AM in our conditions). **(B)** Box plots showing the distributions of mean seizure frequency for dark and light periods of animals treated with saline or PCPZ. *t-test p<0.05. **(C)** Summary plot as in A showing the averaged IID frequency during the daytime, during the test period. Data are normalized to the averaged IID frequency during the control period. **(D)** Same as in B, showing reduced IID frequency in PCPZ-treated mice during both light and dark periods. t-test *p<0.05, ***p<0.001. **(E)** Summary plot as in C, showing the averaged and normalized HFO frequency during the day, during the test period. **(F)** Same as in D, showing normalized averaged HFO frequency. t-test *p<0.05 **p<0.01 ***p<0.001.
